# Molecular basis for the assembly of the dynein transport machinery on microtubules

**DOI:** 10.1101/2024.12.30.630772

**Authors:** Qinhui Rao, Pengxin Chai, Kai Zhang

## Abstract

Cytoplasmic dynein-1, a microtubule-based motor protein, requires dynactin and an adaptor to form the processive dynein-dynactin-adaptor (DDA) complex. The role of microtubules in DDA assembly has been elusive. Here, we reveal detailed structural insights into microtubule-mediated DDA assembly using cryo-electron microscopy. We find that an adaptor-independent dynein-dynactin complex (DD) predominantly forms on microtubules in an intrinsic 2:1 stoichiometry, induced by spontaneous parallelization of dynein upon microtubule binding. Adaptors can squeeze in and exchange within the assembled microtubule-bound DD complex, which is enabled by relative rotations between dynein and dynactin, and further facilitated by dynein light intermediate chains that assist in an adaptor ‘search’ mechanism. Our findings elucidate the dynamic adaptability of the dynein transport machinery, and reveal a new mode for assembly of the motile complex.

## Main Text

Cytoplasmic dynein-1 (dynein) is a motor protein that moves along microtubules (MTs) and plays important roles in a variety of cellular processes, ranging from cargo transport to cell division and the maintenance of cell architecture^1–3^. Dynein is autoinhibited and requires two key cofactors, dynactin and a coiled-coil-containing adaptor, to form the tripartite dynein-dynactin-adaptor (DDA) complex, which is fully activate for processive, unidirectional movement^1,4–6^ (Fig. 1A).

**Fig. 1.**
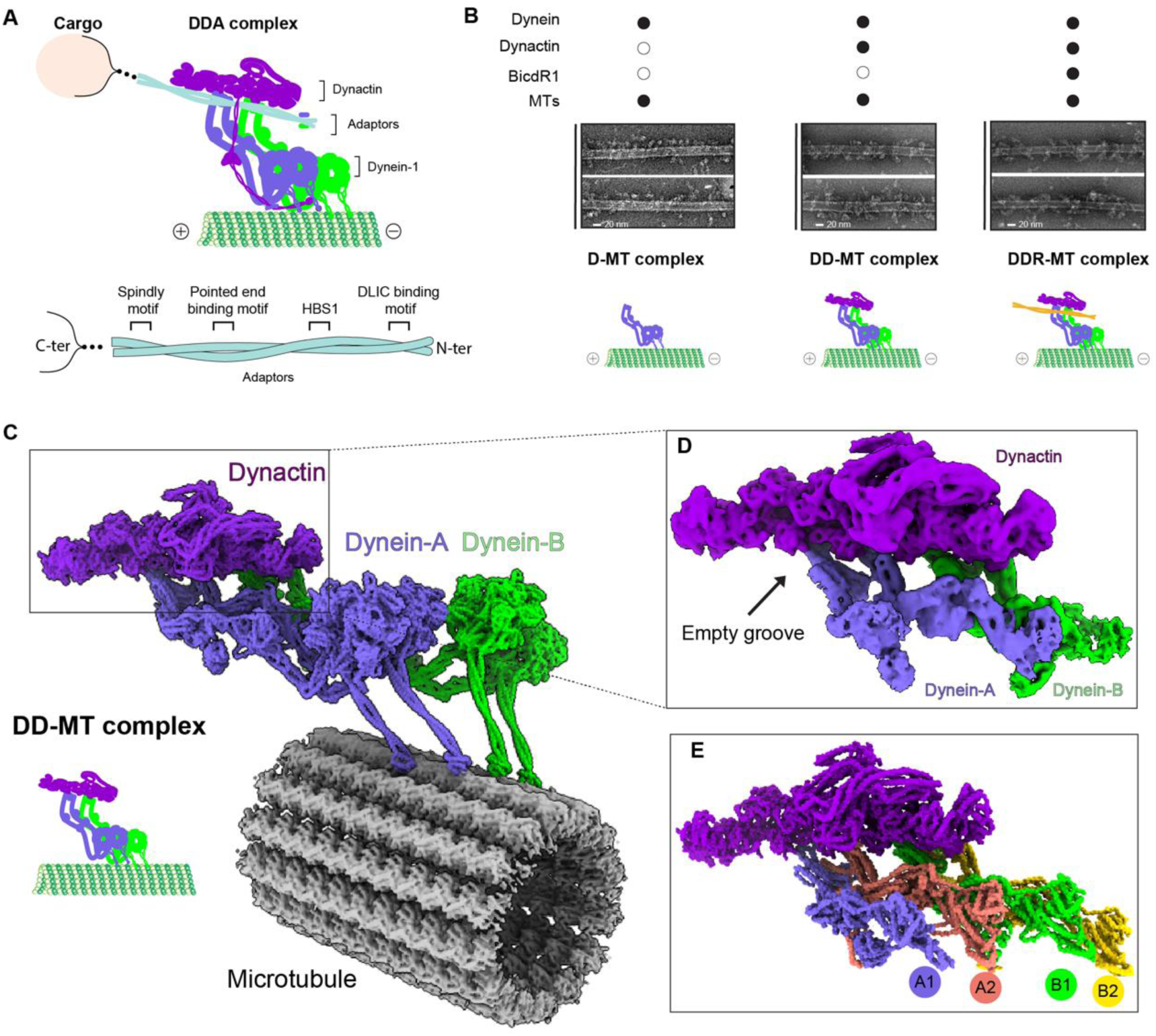
Cryo-EM structure of DD-MT complex. **(A)** Schematic representation of the dynein transport machinery, illustrating dynein, dynactin, and adaptor interactions with cargo (top), and detailed domains of a classical adaptor interacting with dynein and dynactin components, including the spindly motif, pointed end binding motif, HBS1, and DLIC binding motif (bottom). **(B)** Representative micrographs (*n*=50 for each condition) of dynein, dynein-dynactin, and dynein-dynactin-BicdR1 complex bound to MTs in the presence of AMPPNP, with corresponding schematic diagrams depicting the assembly and interaction patterns of these complexes. **(C)** Overall architecture of the DD-MT complex (62,404 particles). **(D)** Density map of the dynein tail interacting with dynactin, with an empty groove indicated. **(E)** Molecular model of the dynein(tail)-dynactin complex, showing dynein (Dynein-A and Dynein-B) and dynactin.

The mechanism of dynein activation is proposed based on studies assessing the assembly of DDA in the absence of MTs, which suggest that the motile complex is assembled prior to landing onto MTs^1,4–8^. It has been suggested that assembly of the motile complex is initiated by the dynein tail domains adopting a parallel alignment, which then permits dynein-dynactin-adaptor binding^4,7^. The conformational signal from the parallel tails is transmitted to the motor domains, triggering them to adopt a parallel conformation, which presumably promotes dynein-microtubule binding and thus unidirectional movement. It has been proposed that the parallel, aligned motors in a DDA complex are indicative of an active, motile conformation. Studies have also shown that the dynein- dynactin binding stoichiometry – i.e., whether dynactin scaffolds one or two dynein dimers – can be determined by the particular adaptor within the tripartite complex^7^. For example, studies indicate that DD-BICD2 complexes mainly scaffold a single dynein, while DD-BicdR1 or DD- HOOK3 complexes recruit two dyneins ^7,9^.

Confounding our understanding of the mechanism of DDA assembly and activation are the somewhat extreme conditions that are required to observe this active complex by structural or single molecule motility methods. For example, adaptors are required to be in large excess (>10 fold) with respect to dynein, and the ionic strength must be kept unnaturally low. Moreover, structural studies have often relied on chemical crosslinking to stabilize the complex, in addition to further purification steps to specifically enrich for assembled complexes^4,7^. Thus, the requisite conditions to assemble DDA *in vitro* are unlikely to be met in a physiological context.

Observations in cultured cells suggest that the dynein transport machinery facilitates long-range transport through an adaptor ‘handoff’ mechanism, which is crucial for autophagosome maturation within primary neuron cells^10^. This model posits that adaptors are interchangeable within a motile DDA complex. Such a mechanism would be inefficient in light of current model, which would first require disassembly of a DDA complex, followed by reassembly with a new cargo adaptor. Moreover, single-molecule motility assays reveal that a significant proportion of dynein molecules are stationary on MTs, with only a subset actively moving^11,12^. These observations raise the possibility that an alternative DDA complex assembly and activation pathway may exist within cells.

MTs are a highly abundant cytoskeletal filament within eukaryotic cells^13^. However, their role in DDA assembly has been largely overlooked in current models. To address their role in dynein activation, we employ a systematic structural investigation by cryo-electron microscopy (cryo-EM) and AI-based structural prediction methods^14,15^, along with biochemical approaches. Our findings indicate that MT binding is sufficient to align the dynein motor domains in the absence of dynactin and an adaptor, and can promote 2:1 dynein-dynactin binding stoichiometry (DD-MT complex), which subsequently serves as a platform for adaptor binding and exchange. Recruitment of adaptors is facilitated by two dynein light intermediate chains (DLICs), each from a different dynein of the DD-MT complex. Dynamic rotations between dynein and dynactin within the DD- MT complex enables efficient adaptor exchange on MTs without the need for disassembly and reassembly of a new DDA complex. Our findings provide an alternative pathway for DDA complex formation mediated by MTs, and substantially expands our understanding of this process within cells.

## Results

### Spontaneous adaptors-independent dynein-dynactin complex formation on MTs

In the absence of MTs, cargo adaptors are required for interactions between the dynein tail domains and dynactin, and thus for processive dynein motility. However, it also has been reported that dynactin alone can enhance dynein run length by approximately 2-fold^16^. We therefore asked if dynactin could interact with dynein on MTs in the absence of adaptors in an attempt to resolve these conflicting findings.

To test this, we purified native dynein and dynactin from pig brains, and performed microtubule pelleting assays (Extended Data Fig. 1A). The results indicate that dynein and dynactin can form a complex on the MTs in the presence of AMPPNP (Extended Data Fig. 1B), with the complex demonstrating high stability up to physiological salt conditions (i.e., 150 mM KCl; Extended Data Fig. 1C). Dynein alone binds well to MTs when it is in a presumably post-powerstroke state (i.e., in the absence of nucleotide, and in the presence of AMPPNP and ADP) (Extended Data Fig. 1D, E), consistent with previous reports for both cytoplasmic dynein-1^17^ and outer-arm dynein (OAD)^18^. Remarkably, dynactin significantly increases the quantity of dynein associated with MTs in all tested nucleotide states, with about 75% of total dynein associated with MTs in the presence of AMPPNP (Extended Data Fig. 1D, 1E). We find the ability of dynactin to enhance MT-binding by dynein also occurs with dynactin obtained from bovine brains, and a recombinant human dynein-1 obtained from insect cells (Extended Data Fig. 1F, 1G). Negative stain electron microscopic examination of these different species revealed that the complex formed by the MT-bound pig dynein and dynactin (DD-MT) closely resembles that of the MT-bound dynein- dynactin-BicdR1 complex (DDR-MT)^19^ (Fig. 1B), suggesting that dynactin enhances dynein-MT binding via interactions with dynein by an adaptor-independent mechanism.

### Intrinsic dynein-dynactin interactions define the 2:1 stoichiometry without an adaptor

Although previous studies have revealed that dynein and dynactin can interact in the absence of cargo adaptors^20–24^, the nature of this complex, and the surfaces that link them together are unclear. For example, it was recently posited that the adaptor-independent dynein-dynactin complex is held together via interactions between the p150 subunit of dynactin and the N-terminus of the dynein intermediate chain (DIC), and that this complex likely does not involve the dynein tail and the ARP1/actin filament of dynactin^22–24^. Current models postulate that stable contacts between the dynein tail and dynactin require a cargo adaptor, and that this is the mechanism by which adaptors promote dynein motility^4,7,19,25,26^. To understand the structural basis for the MT-bound dynein-dynactin complex, we determined its cryo-EM structure (Fig. 1C, Extended Data Fig. 2, Extended Data Table 1, Supplementary Video 1). After 3D classification, we obtained only a single class of dynein-dynactin bound to MTs, which reveals a complex that very closely resembles that of the previously described DDR-MT complex (Extended Data Fig. 2)^19^. In particular, the MT-bound DD complex appears to be stabilized by contacts between the dynein tail domains and the ARP1 filament of dynactin (Fig. 1C-E). Strikingly, we find that this single class for DD-MT, which represents all such complexes, contain two dynein molecules per dynactin, which contrasts with reports indicating that binding stoichiometry is determined by adaptor proteins ^4,7,26^.

Comparison of the DD-MT and DDR-MT complexes shows that the interfaces between the dynein tail and dynactin are almost the same (Extended Data Fig. 3A, 3B). Specifically, the dynein-dynactin interaction in the DD-MT complex is facilitated by the negatively charged surfaces of the ARP1 filament within dynactin and the positively charged surfaces of the dynein tails (Extended Data Fig. 3C-G). These interfaces are reinforced by the complementary geometry between the dynein tail and the ARP1 filament (Extended Data Fig. 4). Helix bundle 1 (HB1) of the dynein tail attaches to one side of ARP1 (actin for A1), while HB2 connects to an adjacent ARP1 (capZ for B2) (Extended Data Fig. 4A-C). Superimposition of all four dynein tails and their bound ARP1s reveals that the dynein tails of chains A2 and B1 align well in the middle, while A1 and B2 rotate away from each other (Extended Data Fig. 4D). This is caused by dynein tail A1 having a slight upward shift, and B2 having a downward shift, which reflects the geometry of the microtubule protofilament angle changes^18,27^.

### Aligned motors may serve as the prerequisite for dynein activation or processivity

To understand how and whether dynactin binding affects the morphology of MT-bound dynein, we used cryo-EM to determine the architecture of dynein alone bound to MTs in the presence of AMPPNP. This revealed that 73.7% of the MT-bound dyneins possess a motor domain with a second poorly visible motor (due to flexibility and heterogeneity of the motors with respect to each other), while the other 26.3% possess two clearly aligned motors (Extended Data Figs. 5 and 6, Extended Data Table 1, Supplementary Video 2). The entire dynein tail in those classes with only one visible motor exhibits too much heterogeneity to be observed clearly (Extended Data Fig. 5A). In contrast, whereas the N-terminal dimerization domain (NDD) and helix bundles 1 (HB1) to 4 (HB4) of the dynein tail in those classes with two aligned motors are dynamic, the remaining regions of dynein tail and liker are sufficiently homogenous that we can identify structural features in our cryo-EM structure (Extended Data Fig. 6).

The structural arrangement of the two aligned motor domains closely resembles that of OAD bound to doublet MTs^18^, suggesting that MT binding can effectively align the dynein motors in the absence of other factors (e.g., dynactin or an adaptor) (Extended Data Fig. 6C). Thus, this arrangement of the motor domains appears to be a universal response of dynein proteins to microtubule binding. Furthermore, the aligned motors may induce a parallel orientation of the tail domains (Extended Data Fig. 6), thus potentially facilitating dynein-dynactin complex formation on MTs. This is the opposite of the current model that posits that dynactin-adaptor binding to the dynein tail facilitates alignment of the motors, which was proposed to be the key event instigating activation of dynein motility. Although the dynein motor domains in both the absence and presence of dynactin can be aligned when bound to MTs, neither exhibits processive motility. Thus, aligned motors may not be an indicator of active dynein, but instead may serve as a prerequisite for dynein activation.

### A dynamic door-opening mechanism for accessibility of adaptors to the pre-assembled DD- MT complex

Our DD-MT structure causes us to reevaluate the role of adaptors in dynein-dynactin interactions. The major interfaces between adaptors and the dynein-dynactin complex are facilitated by electrostatic interactions, with the negatively charged surfaces of the adaptors contacting the positively charged surfaces of the adaptor binding groove within the DD complex (Extended Data Figs. 3C and 7). This electrostatic complementarity likely enhances the affinity and specificity of adaptor binding, enabling efficient formation of the adaptor-DD complex. However, the absence of adaptors in the DD-MT complex raises a key question of whether adaptors could subsequently access and activate the pre-formed dynein-dynactin complex to initiate processivity. In the prevailing model for dynein activation, the formation of an active DDA complex precedes its movement along MTs, while the adaptor binding groove within the DD-MT complex appears to restrict the entry of rod-shaped adaptors due to its geometry (Fig. 1D).

To address this, we first analyzed the structural dynamics of the dynein-dynactin complex on MTs using extensive 3D classification. This reveals that the adaptor-binding groove created by the interaction between the dynein tail and dynactin exhibits considerable dynamics in the absence of adaptors (Fig. 2A, Supplementary Video 3), but is stabilized when the adaptor BicdR1 is bound, as revealed by our cryo-EM structure of DDR-MT, and by that previously reported^19^ (Fig. 2B, Supplementary Video 4). Measurements of the narrow region of this groove – corresponding to the first adaptor binding site (HBS1-A) – indicate that it can accommodate structures with widths between 18.8 to 24.5 Å, closely aligning with the average diameter (∼20 Å) of coiled-coil adaptors (Fig. 2B, C). The second adaptor binding site (HBS1-B) exhibits even larger variation, which may provide an adaptive mechanism for recruiting the second adaptor (Fig. 2C). Taken together, these data suggest that the structural dynamics of the adaptor groove provide a temporal window in which the adaptors can bind to a pre-formed MT-bound dynein-dynactin complex, effectively facilitating the activation for subsequent transport activity.

**Fig. 2.**
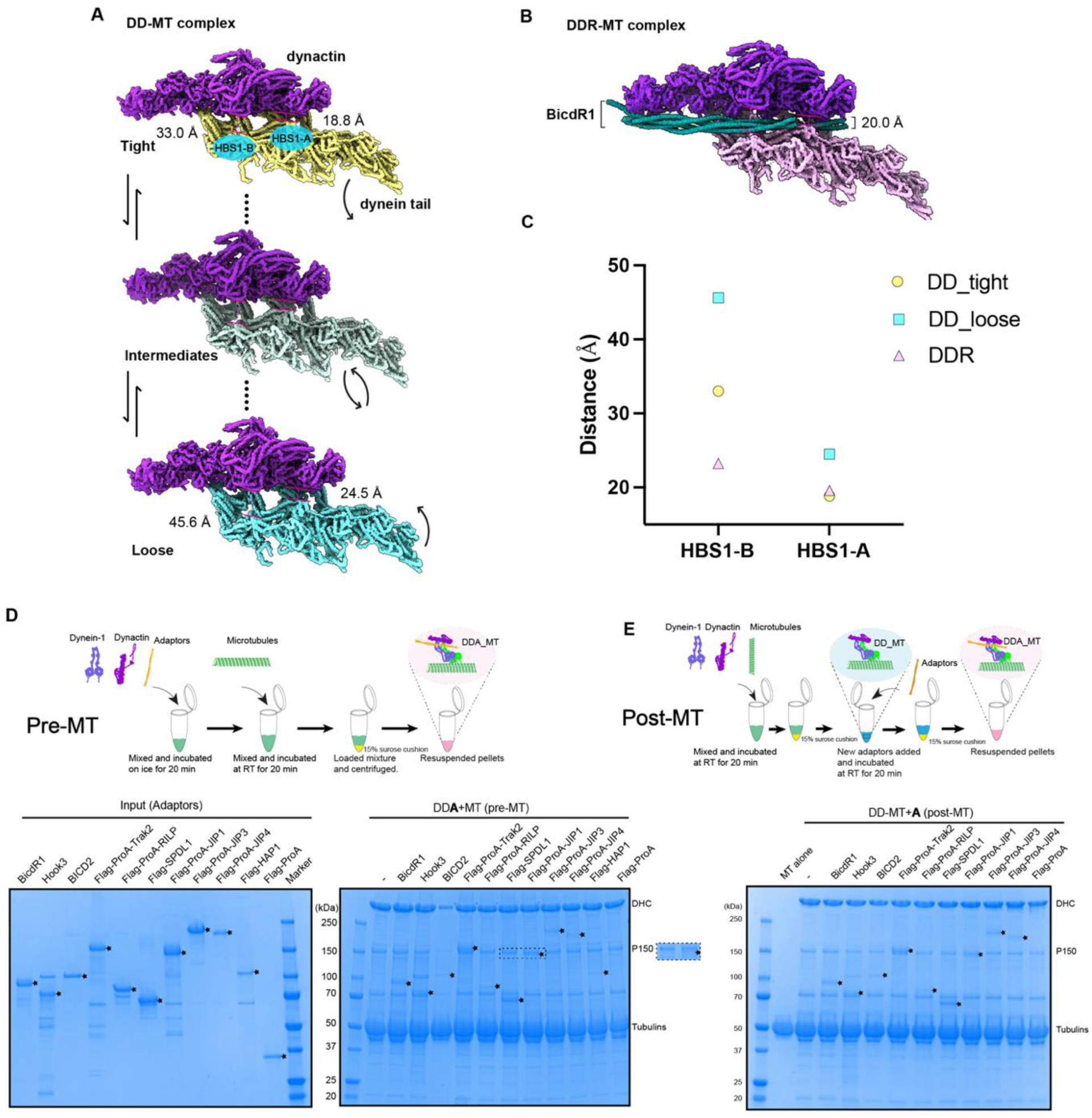
Adaptor-binding groove within the DD-MT complex is dynamically open to accommodate all adaptors. **(A)** A series of dynein-dynactin complex structures showing the dynamic adaptor-binding groove, ranging from tight (top) to loose (bottom). The distances surrounding two HBS1 sites, HBS1-A and HBS1-B, which mediate the interaction between the dynein heavy chain and the HBS1 of adaptors to regulate adaptor entry, were measured. **(B)** Dynamics analysis revealing a stable adaptor-binding groove within the DDR-MT complex. **(C)** Statistical analysis of the HBS1-A and HBS1-B distances in DDR-MT complex, and DD-MT complex in tight and loose states. **(D-E)** Microtubule pelleting assay measuring the binding of dynein-dynactin complex to various adaptors in the pre-MT and the post-MT conditions. Schematic representations of the pre-MT and post-MT processes (top), and SDS-PAGE gels stained with simple blue (bottom). Gels are representative of *n*=3 independent experiments.

To test whether adaptors can bind to DD-MT complexes, we performed microtubule pelleting assays in which we add MTs at different time points (Fig. 2D, 2E). We first mixed dynein and dynactin with one of an assortment of adaptors, incubated the sample for 20 min, and then added MTs prior to pelleting the assembled complex (“pre-MT”). This revealed that all adaptors tested can assemble into an intact DDA complex on MTs (Fig. 2D). We next mixed dynein and dynactin with MTs (to pre-assemble DD-MT complexes), and then added an adaptor prior to pelleting and assessment of complex formation (“post-MT”). This revealed that all adaptors tested are capable of assembling into DDA-MT complexes, and can thus access the adaptor-binding groove of the DD-MT complex, confirming it is indeed accessible to incoming adaptors (Fig. 2E).

### Assembly of dynein transport machinery off MTs is inefficient

We next used negative stain EM to assess the capacity of dynein and dynactin to assemble into DDA complexes with various full-length adaptors in the absence of MTs. Our results reveal that all adaptors except for BicdR1 and HOOK3 fail to interact with dynein and dynactin in these conditions^7^ (Fig. 3A and Extended Data Fig. 8). The formation of the DDA complex in the absence of MTs is thus extremely inefficient, with only ∼3% of the total dynein molecules assembling into dynein-dynactin-BicdR1 (DDR) or dynein-dynactin-HOOK3 (DDH) complexes, in spite of the 10-fold molar excess of BicdR1 or HOOK3 with respect to dynein (Extended Data Fig. 8).

**Fig. 3.**
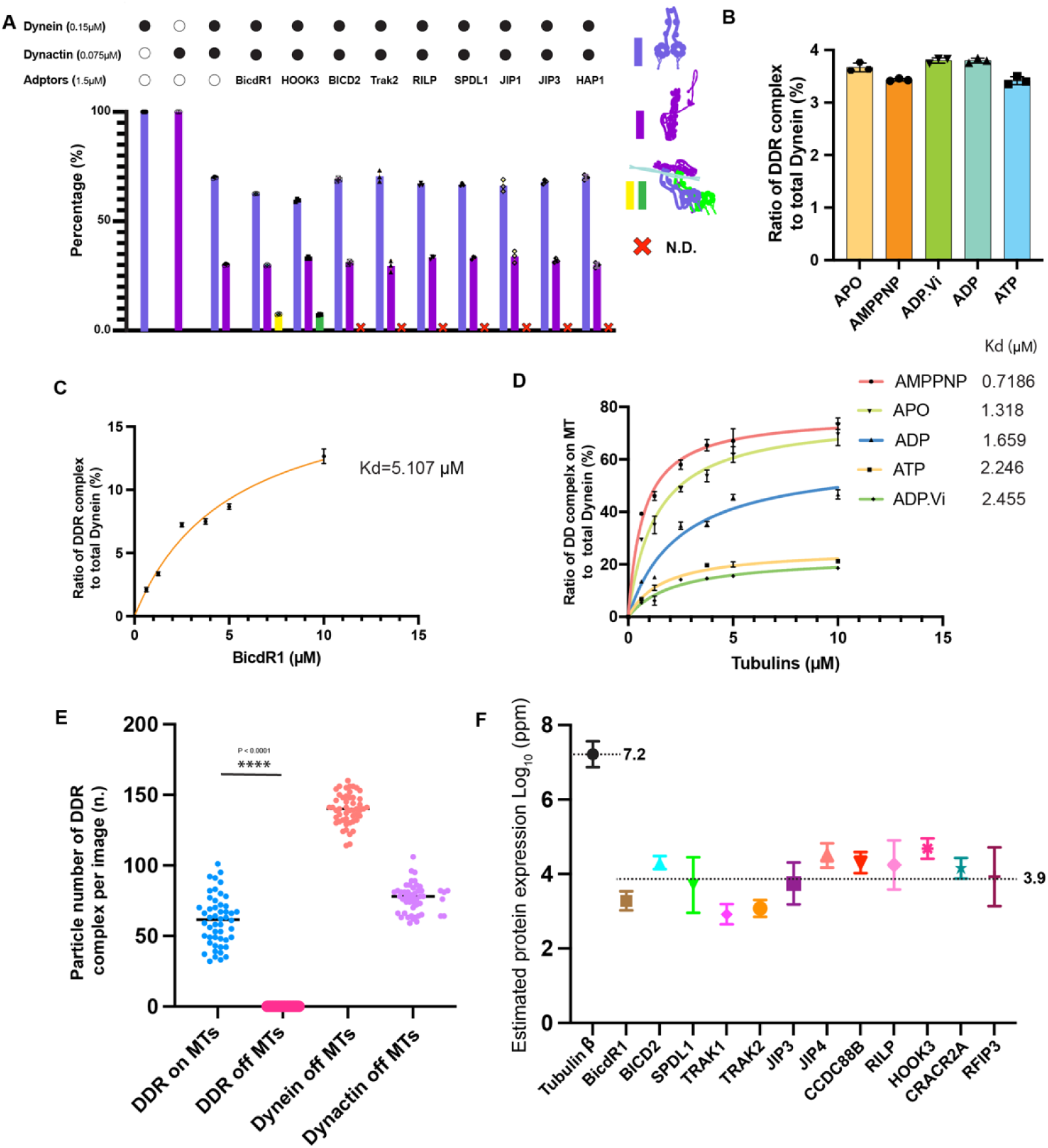
Dynein-dynactin preferentially bind to MTs over adaptor proteins. **(A)** Statistical analysis of interactions between dynein-dynactin and various adaptor proteins (BicdR1, HOOK3, BICD2, Trak2, RILP, SPDL1, JIP1, JIP3, and HAP1) using negative staining. **(B)** Ratio of DDR complex to total dynein in different nucleotide-binding states (Apo, AMPPNP, ADP·Vi, ADP, ATP) using negative staining. **(C)** Binding affinity of dynein-dynactin to BicdR1, with a dissociation constant (Kd) of 5.107 μM measured by using negative staining. **(D)** Measurements of the binding affinity of the dynein-dynactin complex to MTs across various nucleotide-binding states using MT pelleting assay, with the dissociation constants (Kd) listed for each state. **A-D**, Data are mean ± s.d. from *n* = 3 independent experiments. **(E)** Statistical analysis of DDR complex formation on and off microtubules, presented as particle numbers per image (*n*=50 images), analyzed by Mann-Whitney test. **(F)** Average expression levels of tubulin β (*n*=66) and adaptor proteins (BICDR1, *n*=5; BICD2, *n*=48; SPDL1, *n*=13; TRAK1, *n*=16; TRAK2, *n*=25; JIP3, *n*=21; JIP4, *n*=55; CCDC88B, *n*=38; RILP, *n*=26; HOOK3, *n*=57; CRACR2A, *n*=26; RFIP3, *n*=15) in human tissues and cells, data derived from ProteomicsDB, shown in Log_10_ parts per million (ppm) and presented as mean ± s.d..

To understand the low efficiency of DDA assembly off MTs, we predicted the structures of all reported adaptors by AlphaFold^14,15^. We find almost all adaptors are autoinhibited by either blockage of the dynactin and HBS1 binding region by the adaptor’s C-terminus (e.g., BICD2, Spindly)^28–30^, or occupation of the DLIC binding motif by an internal helix (IH) within the adaptor (e.g., Trak2, JIP3)^25^ (Extended Data Fig. 9). In contrast, BicdR1 and HOOK3 exhibit open conformations, explaining their capability of assembling into DDA complexes off MTs. This is consistent with the model that at least some adaptors require an additional uninhibition step (via additional cofactors or cargoes) that thus facilitate the interaction^8,25,29,31^. However, the autoinhibited nature of adaptors likely does not explain the inefficient nature of MT-independent DDA assembly.

### Dynein and dynactin prefer to form DD complexes on MTs prior to adaptor binding

Our data thus far suggest that DD binding to MTs may be preferential to DD binding to adaptors. To address this, we compared the relative binding affinity of the DD complex for adaptors versus MTs. We measured the relative binding affinity of the DD complex for the adaptor BicdR1 in the absence of MTs using a negative stain EM-based approach. Given the relationship between dynein’s nucleotide-bound states and its conformation^32^, we first investigated whether dynein affinity for an adaptor is influenced by different nucleotides. This revealed no significant impact, suggesting that the interaction between the dynein-dynactin complex and adaptors is not influenced by the nucleotide-bound state (Fig. 3B). In the presence of excess BicdR1 (66-fold), the fraction of dynein bound in a DDR complex is ∼12%, with a measured K_d_ of ∼5 µM for BicdR1 to the dynein-dynactin complex (in the presence of AMPPNP; Fig. 3C).

We quantified the binding affinity of the dynein-dynactin complex for MTs in the presence of different nucleotides using microtubule pelleting assays. Our measurements revealed binding affinities ranging from 0.7 µM to 1.6 µM in AMPPNP, ADP, and Apo conditions (states that trigger high MT-binding affinity), with the corresponding binding maxima (Bmax) ranging from 50% to 75% (Fig. 3D and Extended Data Fig. 10). Thus, the apparent affinity of DD for MTs is approximately five times higher than it is for adaptor proteins, indicating a marked preference for microtubule binding over adaptor attachment by the DD complex. Even under ATP and ADP-Vi conditions (conditions that trigger low MT affinity), the binding affinity of the DD complex for MTs is ∼2.3 µM, with a Bmax of ∼20% of total dynein bound to MTs, which is still higher than that for adaptors.

To further verify this, we incubated dynein, dynactin, BicdR1, and MTs in the presence of AMPPNP, then directly performed negative-stain EM analysis without any additional enrichment steps. Along MTs, we observed ∼60 particles corresponding to an intact DDR complex per micrograph (Fig. 3E and Extended Data Fig. 11). However, in spite of observing numerous dynein complexes and dynactin complexes unbound from MTs in our micrographs, we observed only a very small number of intact DDR, consistent with an inefficient in-solution assembly pathway.

Given our observations that autoinhibited adaptors can assemble into a DDA complex in the presence of MTs, but not in their absence, this suggests that MT-binding by the DD complex somehow stimulates the opening of autoinhibited adaptors for subsequent binding (Figs. 2D and 3A). As a major component of the cytoskeleton, the expression level of tubulin is about 2-4 orders of magnitude higher than adaptors (Fig. 3F). Thus, we posit that dynein and dynactin form a DD- MT complex prior to adaptor binding, especially for those adaptors that are autoinhibited.

### DLIC^helix^ facilitates adaptor recruitment for DDA complex formation on MTs

Structural comparisons between the DD-MT and DDR-MT complexes reveal that adaptor stabilization may require two conserved helices (DLIC^helix^) that interact with the middle two dynein heavy chains (of A2 and B1). These two helices are not involved in the DD-MT interaction, suggesting they are not required for assembly of this complex (Extended Data Fig. 3A). However, prior research has underscored the critical function for this conserved helix in dynein transport machinery processivity^33–36^. Building on these insights, we speculate that the interaction between the DLIC^helix^ and the DLIC binding motif of adaptor proteins might be important for the stability of the DDA complex, or for the recruitment of adaptors, thereby enabling subsequent transport.

To test this, we engineered constructs of BicdR1, HOOK3, and Trak2, each devoid of the DLIC binding motif, to assess their binding affinity for the DD-MT complex. Our findings reveal that adaptors lacking the DLIC binding motif show significantly less binding to the DD-MT complex (Fig. 4A-C). Notably, the removal of the IH within Trak2 (see Extended Data Fig. 9B) leads to increased binding to the DD-MT complex (Fig. 4C), suggesting a competition for IH binding between the DLIC^helix^ and the adaptor coiled-coil during DDA complex assembly on MTs. Therefore, we propose that the pair of DLIC helices play a role in capturing the adaptor, and subsequently reorganizing the adaptors for DD binding (Fig. 4D, Supplementary Video 5).

**Figure 4.**
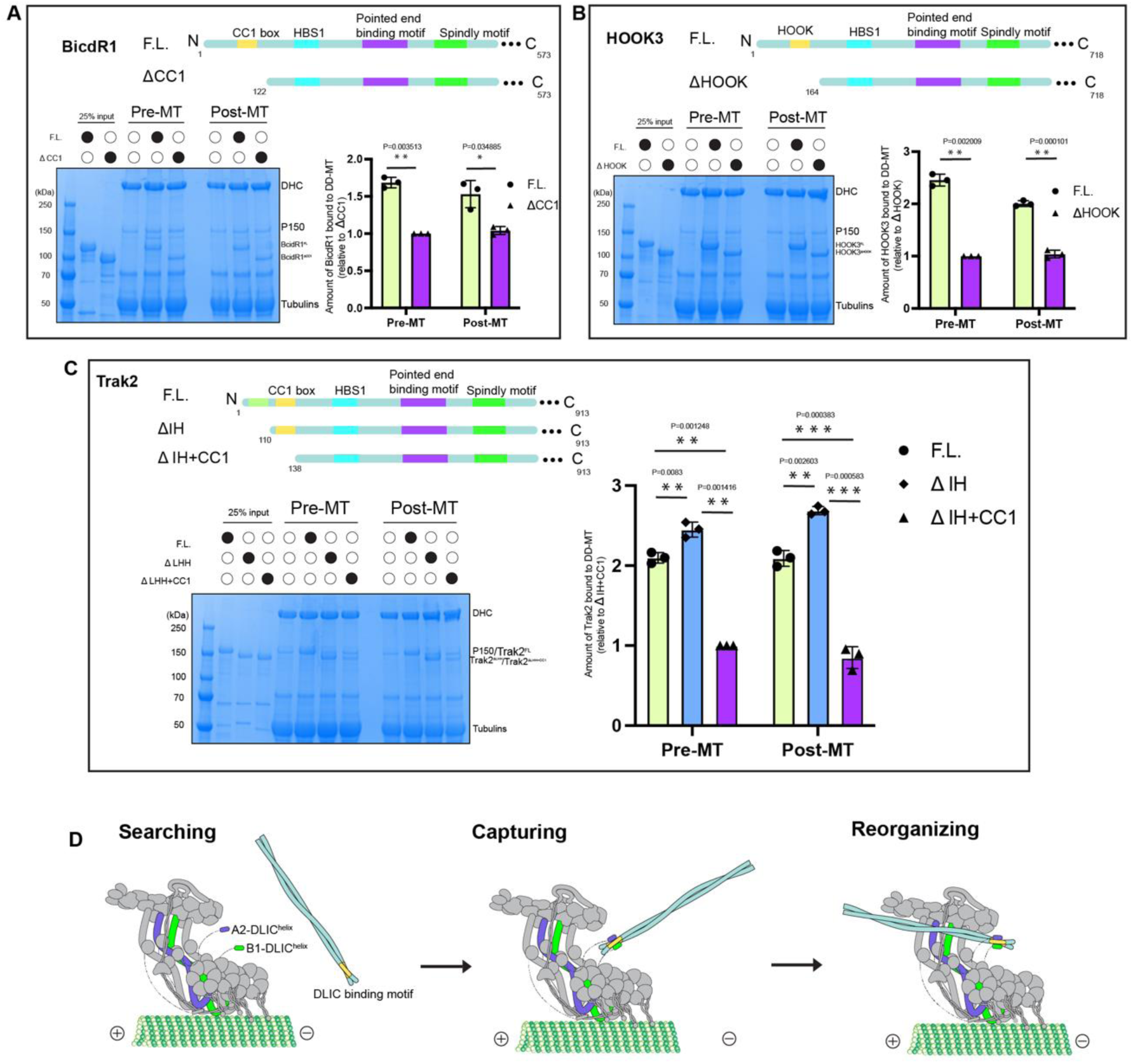
DLIC-binding motif of adaptor is essential for adaptor recruitment. **(A-C)** Schematic representations of BicdR1, HOOK3, and Trak2 adaptors showing their key domains and deletion constructs. Microtubule pelleting assays compare the binding of full-length (F.L.) and deletion constructs (ΔCC1, ΔHOOK, ΔIH, ΔIH+CC1) to the dynein-dynactin complex in pre-MT and post-MT conditions. SDS-PAGE gels stained with simple blue display binding interactions, quantified in bar graphs with statistical significance indicated (right panel). The Y-axis represents the relative amount of adaptor protein compared to each deletion construct: ΔCC1 for BicdR1, ΔHOOK for HOOK3, and ΔIH+CC1 for Trak2. Gels are representative of *n*=3 independent experiments, data are presented as mean ± s.d., and analyzed by unpaired *t*-test with Welch’s correction. **(D)** A proposed model illustrating the stages of adaptor recruitment by the dynein- dynactin complex: searching, capturing, and reorganizing, emphasizing the role of the DLIC- binding motif.

### Adaptor competition and exchange within the DD-MT complex

Based on our observations, we propose that the MT-bound DD complex acts as an initiation complex, performing diffusive migration along MTs and awaiting the arrival of an activating adaptor^6,16^. This suggests potential competition among the various adaptor proteins that may encounter this MT-bound complex in cells. To explore this hypothesis, we conducted an assay to determine whether competition could occur between adaptors Trak2 and BicdR1. Our findings reveal that BicdR1 can efficiently displace pre-assembled Trak2 in the DDK-MT complex (Extended Data Fig. 12A). Conversely, Trak2 is unable to displace pre-assembled BicdR1 in the DD-MT complex (Extended Data Fig. 12A). We also tested a Trak2 variant that lacks the IH domain that blocks the CC1 box binding site (Trak2^ΔIH^). We predicted that deletion of the IH would lead to improved accessibility of the Trak2 CC1 box to DD, which would reduce BicdR1’s ability to compete for DD binding with Trak2. We found this to be the case, as we noted decreased Trak2^ΔIH^ being was competed off by BicdR1 (Extended Data Fig. 12B). Thus, the IH of Trak2, and potentially other adaptors, indeed plays a role in modulating adaptor-DD binding. These data together suggest that adaptors can compete for binding to the MT-bound DD complex, and can exchange with one another.

We sought to confirm the competition between adaptors using cryo-EM. We first assembled the DDK-MT complex and determined its structure. We observed only a single Trak2 dimer within the adaptor-binding groove of the DD complex (Fig. 5A). The density corresponding to Trak2 includes a dynamic and flexible region between the HBS1 and the pointed end binding motif, which is consistent with AlphaFold prediction (Extended Data Fig. 9B). Next, we more closely analyzed our cryo-EM dataset of the DDR-MT complex, and found it consists of two main classes: one possessing two BicdR1 dimers (DDR_2_), accounting for 75% of all complexes, with the remaining 25% possessing a single dimer of BicdR1 (DDR_1_) (Fig. 5B, Extended Data Fig. 13A). We then assembled the DDK-MT complex, and subsequently introduced BicdR1 prior to freezing the samples. Strikingly, nearly all Trak2 adaptors in the MT-bound DDK complexes were replaced with BicdR1, resulting in ∼51% DDR_2_-MT and ∼45% DDR_1_-MT (Fig. 5C, Extended Data Fig. 13B). The remaining ∼4%, termed DDKR-MT, represents an averaged density map containing both BicdR1 and Trak2 (Extended Data Fig. 13B). These results indicate that adaptors can compete and exchange within a DD-MT complex without relying on the disassembly and reassembly of the dynein transport machinery (Fig. 5D).

**Fig. 5.**
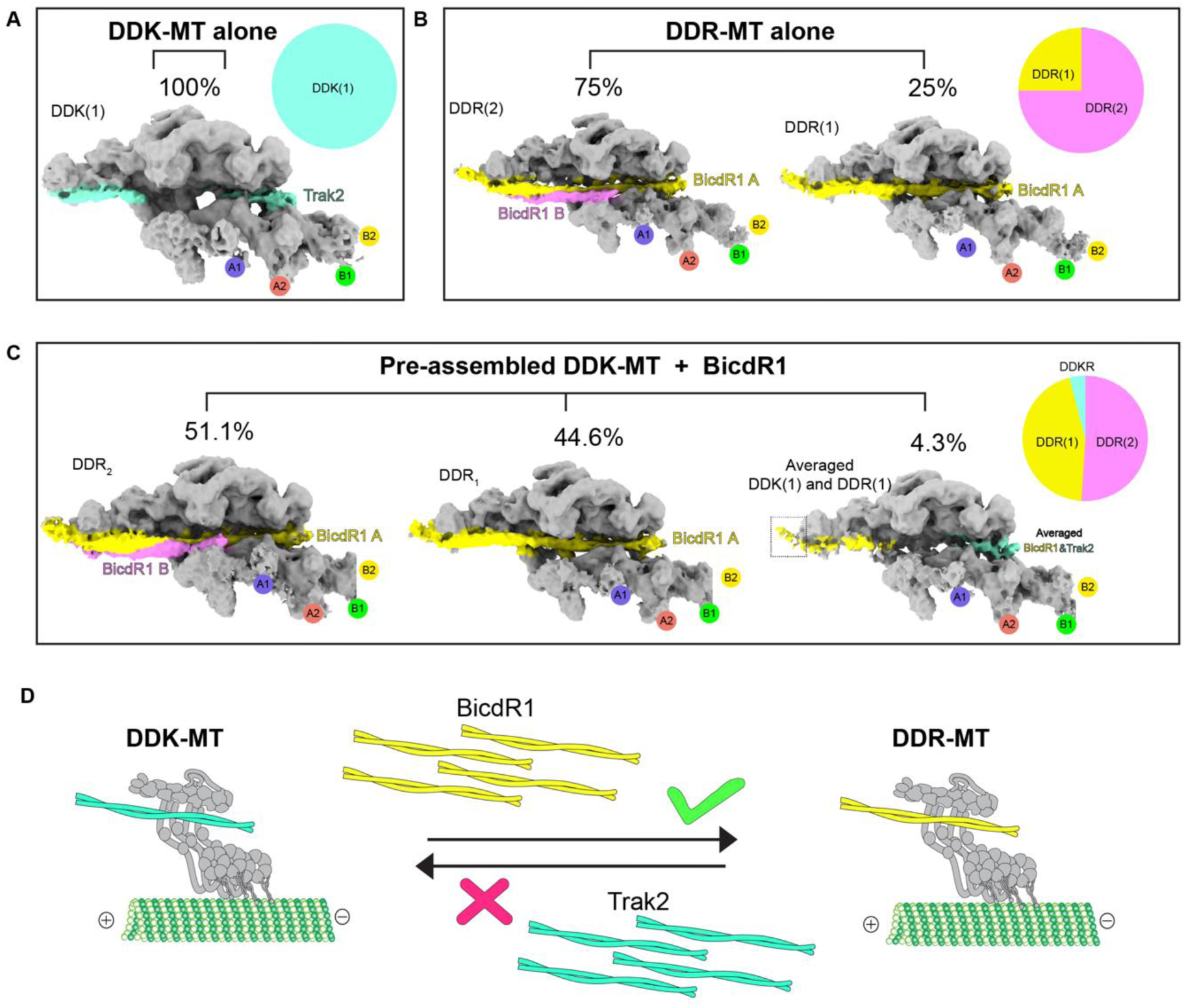
Adaptors compete for binding to the DD-MT complex. **(A)** Density map of the dynein (tail)-dynactin-Trak2 complex (DDK-MT, 52, 920 particles), showing 100% occupancy by Trak2 (steel blue). **(B)** Density maps of the dynein (tail)-dynactin-BicdR1 complex (DDR-MT, 42, 951 particles), showing a distribution of 75% DDR_2_ and 25% DDR_1_. BicdR1-A and BicdR1-B are colored by yellow and magenta, respectively. DDR_2_ and DDR_1_ indicate two and one dimer of BicdR1 within DDR complex, respectively. **(C)** Density maps illustrating the competition results that excess BicdR1 incubating with the pre-assembled DDK-MT complex. The resulting binding distribution is 51.1% DDR_2_ (33, 721 particles), 44.6% DDR_1_ (29, 431 Particles), and 4.3% averaged binding of DDK_1_ and DDR_1_ (2, 837 Particles). The pie chart summarizes the proportions of these complexes. **(D)** Schematic representation of adaptor competition, showing that BicdR1 can compete Trak2 for binding to the dynein-dynactin complex.

## Discussion

### A revised model for assembly of the dynein transport machinery

Our research identifies a new function for MTs in assembly and activation of the dynein transport machinery, leading us to propose an updated model that underscores their significant influence (Fig. 6). Our findings demonstrate that the binding of dynein to MTs results in the alignment of the two motor domains. We hypothesize that this somehow triggers the tail to adopt a parallel orientation, which consequently promotes efficient dynactin binding (Fig. 6A-C), and also enables the recruitment of a second dynein dimer to the MT-bound complex independent of adaptor proteins (Fig. 6D). Thus, the DD-MT complex likely serves as a pre-initiation complex, which is yet to be activated (Fig. 6D). Subsequently, the complex utilizes two DLIC^helices^ (within the two middle dynein complexes) to assist in the ‘search’ for a potential adaptor (Fig. 6E). For long-range transport of cargos that require multiple adaptors^10^, the incoming adaptor with a higher binding affinity can replace the current adaptor, facilitating continuous transport without the need for disassembly and reassembly of the dynein transport machinery (Fig. 6E-F).

**Fig. 6.**
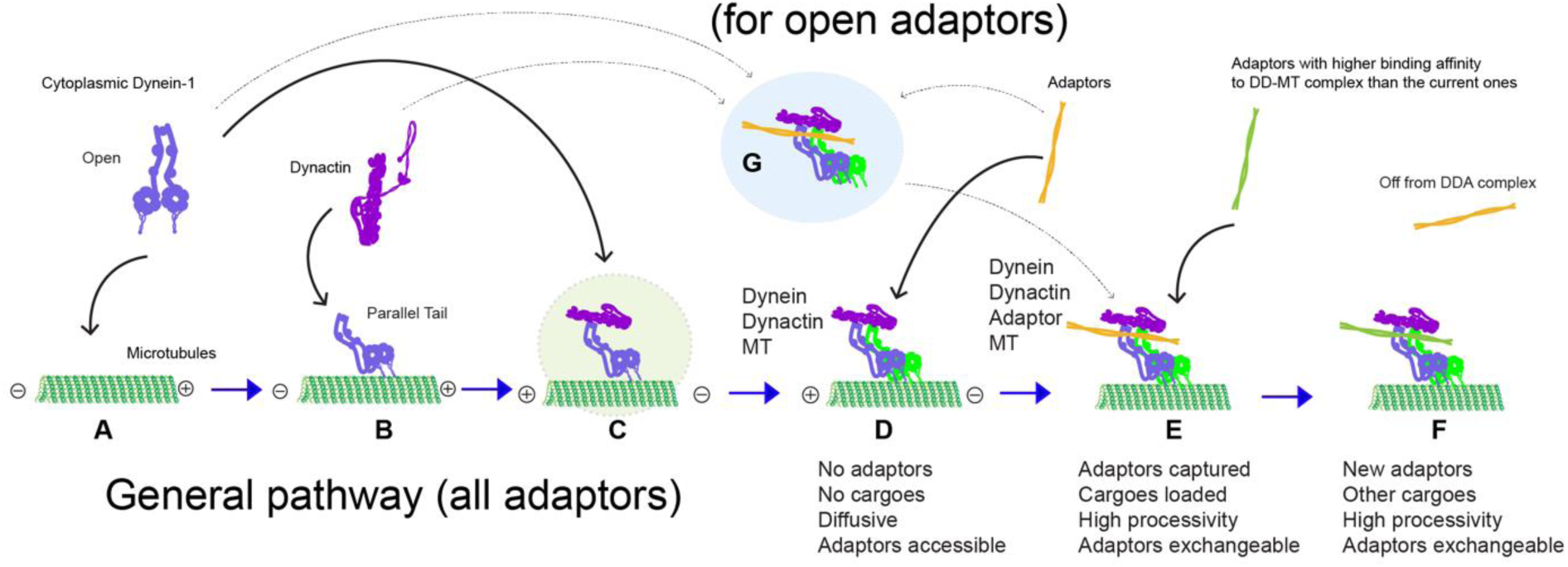
Model illustrating dynein transport machinery assembly and adaptor binding. General pathway for all adaptors (A-F, this study, black and blue arrows): **(A)** MT is available to bind open dynein transitioning from a phi-particle state. **(B)** Dynein adopts a parallel tail conformation upon binding to MTs. **(C)** Dynactin is recruited, forming a pseudo-intermediate DD- MT complex. **(D)** Fully assembled DD-MT complex with a molar ratio of 2:1, capable of recruiting adaptors for cargo transport. **(E)** Adaptors with higher binding affinity compete for binding to DD- MT, replacing current adaptors. **(F)** DDA complex with new adaptor assembled on MT for new cargo transport. **Pathway for open adaptors (G to E, previous studies**, **dashed arrows)**: **(G)** DDA complex forms off MTs and becomes fully activated after MT binding. Blue arrows indicate DDA-MT complex formation in the general pathway.

The prevailing model posits that an activated DDA complex first assembles in the cytoplasm, and then lands on MTs to initiate cargo transport^1,2,4,25^ (Fig. 6G). Our findings suggest that the formation of the DDA complex off MTs is a minor pathway. Although a DDA complex can be assembled in the cytoplasm with help from cofactors, such as LIS1^1,25,37^, its delivery to MTs (via diffusive movement throughout the crowded cytoplasm) is likely an inefficient process due to its enormous size^1,38^. In contrast, dynein-dynactin exhibits a significantly higher affinity for MTs than for adaptors (Fig. 3C, D). Moreover, the adaptors’ expression levels and consequent abundance are much lower than tubulin (Fig. 3F). Thus, binding of dynein and dynactin to MTs is a more likely scenario, which facilitates the assembly of motile DDA complexes that are even competent for adaptor exchange. Our results strongly indicate that this is a major pathway for DDA complex assembly. Moreover, the ability of MT-bound DD to undergo adaptor exchange likely facilitates long-range transport of cargos (e.g., autophagosomes) that require multiple adaptors (Fig. 6A-F).

### MT-bound dynein-dynactin is a pre-initiation complex

Recent single molecule studies have shown that the adaptor-free dynein-dynactin complex exhibits diffusive or stationary behavior on MTs, indicating that the DD complex is inactive, and requires adaptors for activation^6^. Single-molecule imaging of DD in live cells has also shown that these two complexes colocalize on MTs, and are predominantly in a paused state on MTs during retrograde transport, suggesting that most of the dynein transport machinery is inactive in vivo^11,12^. This allows us to speculate that the DD-MT complex exists in a paused, adaptor-free state on MTs in cells, and is awaiting adaptor-bound cargo binding to initiate transport (Fig. 6D). This idea is supported by studies in fission yeast which suggest that dynein first binds and diffuses along MTs prior to binding to the presumed adaptor protein Mcp5, which stimulates minus-end movement^39^.

### One dynactin intrinsically scaffolds two dyneins

Previous studies have indicated that dynein activity can be modulated by various adaptors. Different dynein-dynactin binding stoichiometries, which can be dictated by specific adaptors, have been correlated with differing degrees of dynein activity (e.g., velocity, force generation)^7,40^. For example, those DDA complexes mostly comprised of 1 dynein:1 dynactin:1 adaptor (e.g., DD- BicD2) exhibit lower velocities than those comprised of mostly 2:1:1 (DD-BicdR1 or DD- HOOK3)^4,7^. Of note, previous studies have found that the inclusion of MTs can shift the ratios of DD-BicD2 to 2:1:1 (*9*), and even promotes stable recruitment of a 2^nd^ cargo adaptor. Integrating previous findings with our research, we’ve confirmed a consistent dynein to dynactin ratio of 2:1, regardless of adaptor identity. This allows us to conclude that the binding stoichiometry of dynein to dynactin – at least in the presence of MTs – occurs by an adaptor-independent mechanism, possibly due to the tails adopting a parallel configuration.

Our comprehensive 3D classification reveals that dynein alone exhibits various binding patterns to MTs, including a significant proportion (26%) in which the two motor domains are aligned in a parallel configuration (Extended Data Fig. 5). Notably, patterns of four aligned motors, as might be expected in a DDA-MT complex, were not observed in the absence of dynactin. This absence suggests that dynein’s attachment to MTs occurs in a stochastic manner. Upon introducing dynactin, dynein’s binding patterns were consistently observed to involve two dyneins after the extensive 3D classification, in which motors of dynein-B is always in aligned, and motors of dynein-A can be either in aligned or staggered (Extended Data Fig. 2A). This observation leads us to speculate that dynactin may play a crucial role in orchestrating the recruitment and subsequent arrangement of the second dynein within a 2:1 DD-MT complex.

### Adaptor recruitment and exchange is regulated by DLIC^helix^

Our analysis suggests that recruiting two dynein molecules is crucial for adaptor binding. The interplay between the two DLIC^helices^ (on dynein A2 and B1) may be key in competing with the IH within some adaptors (e.g., Trak2), thus relieving their autoinhibited state and enabling their binding to the DD-MT complex. This competition necessitates the simultaneous presence of the two LIC helices, explaining why the dynein-dynactin complex can efficiently recruit adaptors in the presence of MTs but struggles in their absence, especially for autoinhibited adaptors. This also helps to explain in part why assembly of DDA complexes in solution is inefficient.

We propose that the initial recruitment of the first adaptor is fundamentally reliant on the presence of these two DLIC^helices^. The subsequent incorporation of a second adaptor appears to be easiest for BicdR1, an adaptor that is not autoinhibited. For those adaptors that are autoinhibited, a single DLIC^helix^ from heavy chain A1 may struggle to effectively compete for binding with the DLIC binding motif in the presence of the adaptor^IH^, potentially resulting in the absence of a second adaptor, as noted with DDK-MT and DDJL-MT^25^. Even for uninhibited adaptors, there is only a subset of DD-MT complexes that possess a second adaptor (Fig. 5B, C). Finally, the exchange of adaptors may begin with the competition between the DLIC binding motifs of the new adaptor and the currently bound adaptor, along with the two DLIC^helices^. If the new adaptor has a higher binding affinity for the DLIC^helices^, then the replacement can occur efficiently; otherwise, the exchange may fail (Fig. 5D). It is important to note that other binding interfaces between adaptors and the dynein-dynactin complex are also involved, and thus the mechanism of adaptor exchange will require further investigation.

## Methods

### Dynein and Dynactin purification

The purification of endogenous dynein and dynactin from pig brain involved a series of precise and systematic steps to isolate these protein complexes with high purity, as previously described41. Initially, 500 grams of frozen pig brain tissue was processed through mashing and subsequent solubilization in a homogenization buffer containing 35 mM PIPES (pH 7.2), 5 mM MgSO4, 1 mM EGTA, 0.5 mM EDTA, 0.4 mM ATP, 1x protease inhibitor cocktail, 1 mM PMSF, and 1 mM DTT. This resulted in a homogenized brain mixture, which was first clarified by centrifugation at 16,000 rpm for 20 minutes. The resulting supernatant was then subjected to a second centrifugation at 140,000 g for 1 hour to obtain a cleaner supernatant. This was subsequently filtered through a nylon filter with 40 μm pores to remove any remaining particulate matter.

The filtered supernatant was applied to a pre-equilibrated SP-Sepharose column with homogenization buffer, where it was washed twice to remove non-specifically bound material. Dynein and dynactin proteins were then eluted from the column using homogenization buffer supplemented with 0.6 M KCl. The fractions containing the peak protein were collected for further purification.

To enrich the dynein and dynactin complexes, the peak fractions were subjected to density gradient centrifugation. This was achieved by layering the fractions over a sucrose cushion consisting of 60% and 20% sucrose layers and centrifuging at 140,000 g for 11 hours. The fractions containing dynein and dynactin complexes were carefully collected and reloaded onto an SP-Sepharose column for a second round of purification, ensuring the removal of residual contaminants.

For further refinement, the eluted fractions were applied to a gradient ranging from 10% to 40% sucrose and centrifuged at 140,000 g for 17 hours. The fractions containing dynein and dynactin proteins were monitored using SDS-PAGE, after which they were applied to a pre-equilibrated Mono Q column with a Mono Q buffer. The buffers used were Buffer A (20 mM Tris, pH 7.2, 30 mM KCl, 1 mM MgCl2, 1 mM DTT, 0.5 mM ATP) and Buffer B (20 mM Tris, pH 7.2, 1 M KCl, 1 mM MgCl2, 1 mM DTT, 0.5 mM ATP). The dynein complex and dynactin were eluted from the Mono Q column using linear gradients of Buffer B, tailored to each protein complex’s specific binding and elution characteristics. These fractions were collected, analyzed by SDS-PAGE for purity assessment, and further characterized by negative staining to confirm the integrity and composition of the purified complexes.

### Adaptors expression and purification

Full-length BicdR1 and HOOK3 proteins were produced in insect cells and purified following established protocols7. Full-length BICD2 was expressed in BL21 E. coli cells and underwent a purification process as previously detailed28. Plasmids encoding the full-length versions of JIP1, JIP3, JIP4, RILP, and Trak2 were acquired from Addgene. Briefly, pCDNA3 T7 Jip1 was a gift from Roger Davis (Addgene plasmid # 51699)42; GFP-JIP3/4 was a gift from Mark Cookson (Addgene plasmid # 164624, Addgene plasmid # 164620)43; pTRE2-Bla(HA–RILP-FLAG) was a gift from Steven Weinman (Addgene plasmid # 102424)44; and GFP-Trak2 was a gift from Josef Kittler (Addgene plasmid # 127622)45. Meanwhile, the genes encoding HAP1 and Spindly were synthesized by Twist Bioscience. These genes were then inserted into a mammalian expression vector, specifically a pCAG-Flag-ProteinA (or pCAG-Flag-MBP) plasmid, designed to facilitate protein expression and purification.

Following the cloning process, the constructs were introduced into Expi293 cells—a cell line optimized for high-efficiency transfection and protein production—via transfection. The expressed adaptor proteins were then isolated using anti-Flag agarose gel, leveraging the affinity of the Flag tag for a straightforward purification process. Subsequently, the proteins were eluted from the gel using a 3xFlag peptide, which competes with the bound protein for binding sites on the agarose gel, effectively releasing the protein. All adaptor protein were collected, concentrated, and stored in a buffer containing 25 mM HEPES pH 7.4, 150 mM NaCl, 1 mM DTT, 10 % glycerol.

### Microtubule pelleting assay

To conduct the pelleting assay, a 20 μL reaction system was meticulously prepared. Initially, 15 μL of dynein and associated complexes were mixed thoroughly on ice for 15 minutes to form a pre-incubation mixture. This mixture was then allowed to equilibrate to room temperature (RT) over 5 minutes, ensuring a gradual transition to prevent any thermal shock that could affect the protein’s activity. Following this, a microtubule mixture, comprising 5 μM tubulin and 20 μM Taxol, was gently added and mix. The combined mixture was incubated at RT for an additional 15 minutes to facilitate the binding of dynein or dynein associated complex to the MTs.

To efficiently distinguish between microtubule-bound and unbound components, the reaction mixture was layered over a cushion solution containing 15% sucrose in a specific buffer. This step aids in the separation process during centrifugation. The sample was then centrifuged at 20,000 g for 8 minutes, a condition optimized to pellet the microtubule-protein complexes effectively while leaving unbound proteins in the supernatant. After centrifugation, the supernatant was carefully removed, and the pellet, containing the microtubule-bound fraction, was washed twice with a wash buffer to remove any non-specifically bound proteins.

Finally, the pellet was resuspended in a suitable resuspension buffer tailored for subsequent analyses. This step is crucial for preparing the pellet for downstream applications, whether for biochemical assays or structural studies, ensuring that the microtubule-bound proteins are in an optimal state for further examination.

### Adaptors competing assay

After completing the microtubule pelleting assay, the first adaptor assembled DDA complex is formed and purified using a sucrose cushion. Subsequently, the DDA complex is resuspended and incubated with an excess amount of a second adaptor at room temperature (RT) for 20 minutes before subjecting it to a second sucrose cushion. Both the supernatant and pellet are then collected separately and analyzed by SDS-PAGE.

### D-MT and DD-MT samples preparation

The preparation D-MT and DD-MT samples were followed by microtubule pelleting assay with a scale-up volume and resuspend with a purposed volume for cryo-EM analysis.

### Cryo-EM data collection

All cryo-EM grids were screened at the Yale ScienceHill-Cryo-EM facility using a Glacios microscope (Thermo Fisher Scientific). Subsequent cryo-EM data were collected at two different sites: Yale ScienceHill-Cryo-EM facility with a Glacios microscope operated at 300 keV with a K3 detector; and the Laboratory for BioMolecular Structure at BNL with a Krios microscope operated at 300 keV with a K3 detector and a Bioquantum Energy Filter. Automatic data collection was facilitated by either SerialEM46 or EPU software. In total, 6110, 10660, 3000, and 8369 movies were acquired for dynein-MT, dynein-dynactin-MT, dynein-dynactin-BICDR1-MT, and dynein-dynactin-TRAK2-MT, respectively. Detailed data collection parameters can be found in Extended Data Table 1.

### Cryo-EM image processing of dynein-MT dataset

Preprocessing steps, including motion correction, CTF estimation, and particle picking, were conducted either in cryoSPARC Live^47^ or via an in-house script utilizing MotionCor2^48^, GCTF^49^, and Gautomatch. Cryo-EM scripts for real-time data transfer and on-the-fly preprocessing are available for download at https://github.com/JackZhang-Lab.

For cryo-EM image processing of dynein-MT dataset, we employed a pipeline previously developed for MT tracing and MT-signal subtraction (Extended Data Fig. 5)^50,51^. Initially, MT particles were detected using template matching with a distance cut-off of 8 nm, with a relatively low cross-correlation score set to include MT particles with low contrast or located at cross-over locations. Following three rounds of 2D classification, false-picked MT particles, including carbon edges, and mis-centered MTs were rejected. The selected coordinates were recentered and subjected to "multi-curve fitting". After MT-signal subtraction, original micrographs were replaced by MT-signal subtracted micrographs.

Subsequently, the blob-picker was used to detect motor domains in these new micrographs. 2D classification was utilized to remove junk particles and residual MT particles, after which the selected particles underwent ab initio reconstruction. All original particles were used for initial heterogeneous refinement. After several rounds of 3D and 2D sorting, classes with clear motor features were selected and subjected to local refinement, global, and local CTF refinement.

For the reconstruction of full-length dynein bound to MT, particles were reextracted using a large box size covering two motor domains. The particles were directly reconstructed into a volume featuring the weak density of the second motor. Subsequently, this map was low-pass filtered to 60 Å for heterogeneous refinement, where some classes exhibited enhanced density for the second motor. We then manually fitted the motor map into the weak density to create a two-motor map. This map, together with previous reconstructions, was used for another round of heterogeneous refinement. Ultimately, three main states were predominantly observed for each dynein-MT dataset: i) two stable and parallel heads, ii) one stable leading head, and iii) one stable trailing head.

### Cryo-EM image processing of dynein-dynactin-MT dataset

For cryo-EM image processing of dynein-dynactin-MT dataset, the preprocessing steps are the same as dynein-MT dataset. For particle picking, a previous published dynein-tail and dynactin map (EMD-4177^7^) was projected and used for template picking. The picked particles were directly subjected into heterogeneous refinement with 8 classes (Extended Data Fig. 2). The class with clear dynactin and dynein tail density was selected for rounds of reference-free 2D classification. High quality particles were selected for high resolution reconstruction.

To reconstruct the dynein motor region, particles were recenter on the motor region and re-extracted. After 3D reconstruction and low-pass filtering to 30Å, the 4 motor domains were clearly visible but with smeared density. Rounds of heterogeneous refinement were used to improve the motor domain and two major classes were identified: aligned and staggered as previously reported^19^. Composite dynein-dynactin map was generated in ChimeraX^52^ by merging local refined maps. Dynein-dynactin-adaptors datasets were processed similarly.

### Model building and refinement

For model building and refinement, we utilized two previously reported dynein and dynactin structures (PDB: 7z8f^19^, 6znl^53^) as initial models. Local regions were rigidly docked into the cryo-EM map in UCSF ChimeraX. Subsequently, Namdinator^54^, a molecular dynamics flexible fitting tool, was employed to further refine the model into the cryo-EM map. Manual inspection and adjustments of the model were carried out in COOT v0.9.5^55,56^.

All models underwent iterative refinement using Phenix real-space refinement 1.21rc1_5190 and manual rebuilding in COOT. The quality of the refined models was assessed using MolProbity^57^ integrated into Phenix^58^, with statistics reported in table S1.

## Acknowledgments

We are very grateful to members of the Zhang lab and Steven Markus for their valuable discussions. We would like to thank S. Wu, K. Zhou and J. Lin for their help with cryo-EM data collection at Yale Cryo-EM facility. We thank L. Wang, J. Kaminsky, and G. Hu at the Laboratory for BioMolecular Structure (LBMS) for help with cryo-EM data collection.

## Funding

National Institutes of Health grant R35GM142959 (KZ)

Pittsburgh Center for HIV Protein Interactions grant U54AI170791 (KZ).

National Institutes of Health grant S10OD023603 (FS)

DOE Office of Biological and Environmental Research KP1607011 (LBMS).

## Author contributions

Conceptualization: QR, KZ Methodology: QR, PC, KZ Investigation: QR, PC, KZ Visualization: QR, PC Funding acquisition: KZ Project administration: KZ Supervision: KZ Writing – original draft: QR Writing – review & editing: QR, PC, KZ

## Data availability

All atomic coordinates and cryo-EM maps have been deposited in the Protein Data Bank (PDB) and Electron Microscopy Data Bank (EMDB) under accession codes 9DGQ/46844 (D-MT, overall structure), 9DGP/46843 (D-MT, dynein motor domain), 9DGR/46845/ (composite DD-MT structure), 9DGS/46846 ( dynactin-dynein tail of DD-MT complex), 9DGT/46847 (DDR1-MT, dynactin-dynein tail-one dimeric BicdR1), 9DGU/46848 (DDR2-MT, dynactin-dynein tail-two dimeric BicdR1), 9DGV/46849 (DDK-MT, dynactin-dynein tail-one dimeric Trak2).

---

**Extended Data Figure 1.**
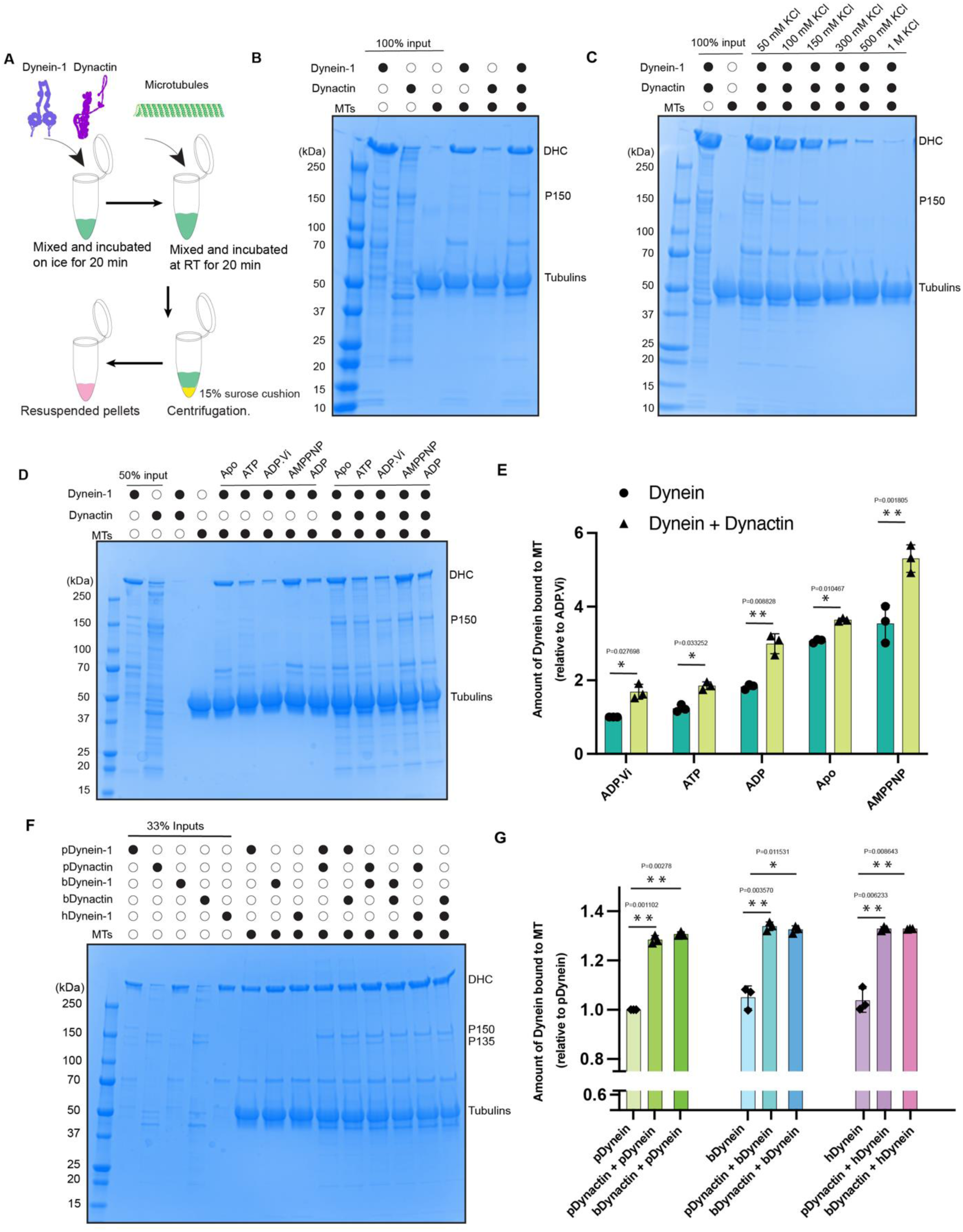
Microtubule pelleting assay reveal the formation of the DD-MT complex. **(A)** Diagram of the microtubule pelleting assay procedure. **(B)** SDS-PAGE gel of dynein, dynactin, and dynein-dynactin complex bound to MTs in the presence of AMPPNP. **(C)** SDS-PAGE gel of dynein-dynactin complex bound to MTs under the salt concentration ranging from 50 mM to 1M in the presence of AMPPNP. **(D)** SDS-PAGE gel of dynein and dynein-dynactin complex bound to MTs in different nucleotide binding states. **(E)** Statistical analysis of the amount of dynein bound to MTs relative to dynein alone bound to MTs in the presence of ADP·Vi. Data are presented as mean ±s.d., analyzed by unpaired *t*-test with Welch’s correction. **(F)** SDS-PAGE gel showing dynein and dynein-dynactin complex from different species (pig, bovine, human) bound to MTs in the presence of AMPPNP. **(G)** Statistical analysis of the amount of dynein bound to MTs relative to dynein alone bound to MTs. Data are presented as mean ± s.d., analyzed by unpaired *t*-test with Welch’s correction. (p, pig; b, bovine; h, human). All SDS-PAGE gels are stained with simple blue. **B, C, D, F,** Gels are representative of *n*=3 independent experiments.

**Extended Data Figure 2.**
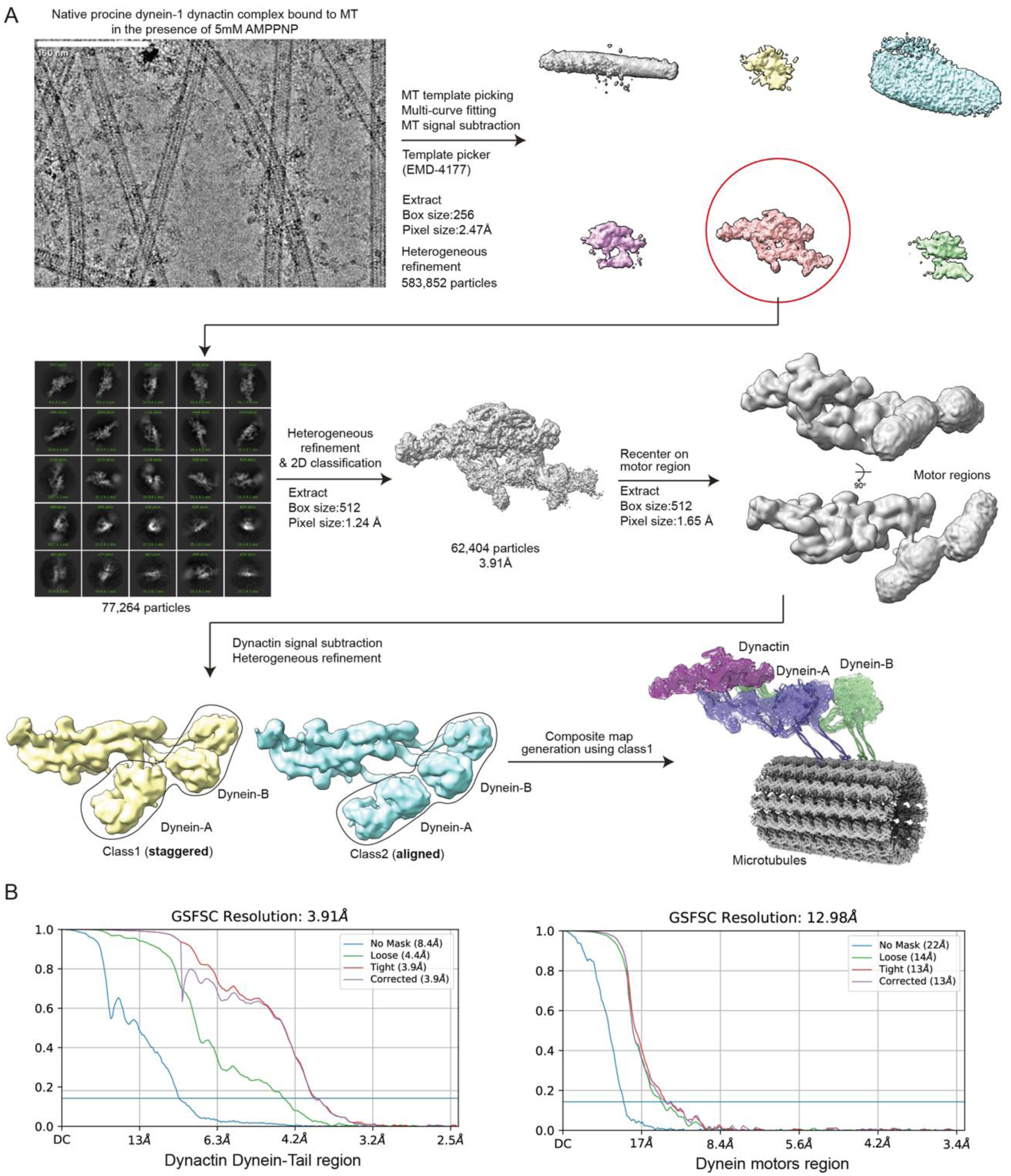
Data processing flow chart of DD-MT. **(A)** Representative image of dynein-dynactin bound to MTs and workflow of cryo-EM image processing. **(B)** FSC curves of dynein tail and dynactin region, and dynein motors region reconstruction. Dynein-dynactin with adaptors (BicdR1, Trak2) datasets were processed similarly.

**Extended Data Figure 3.**
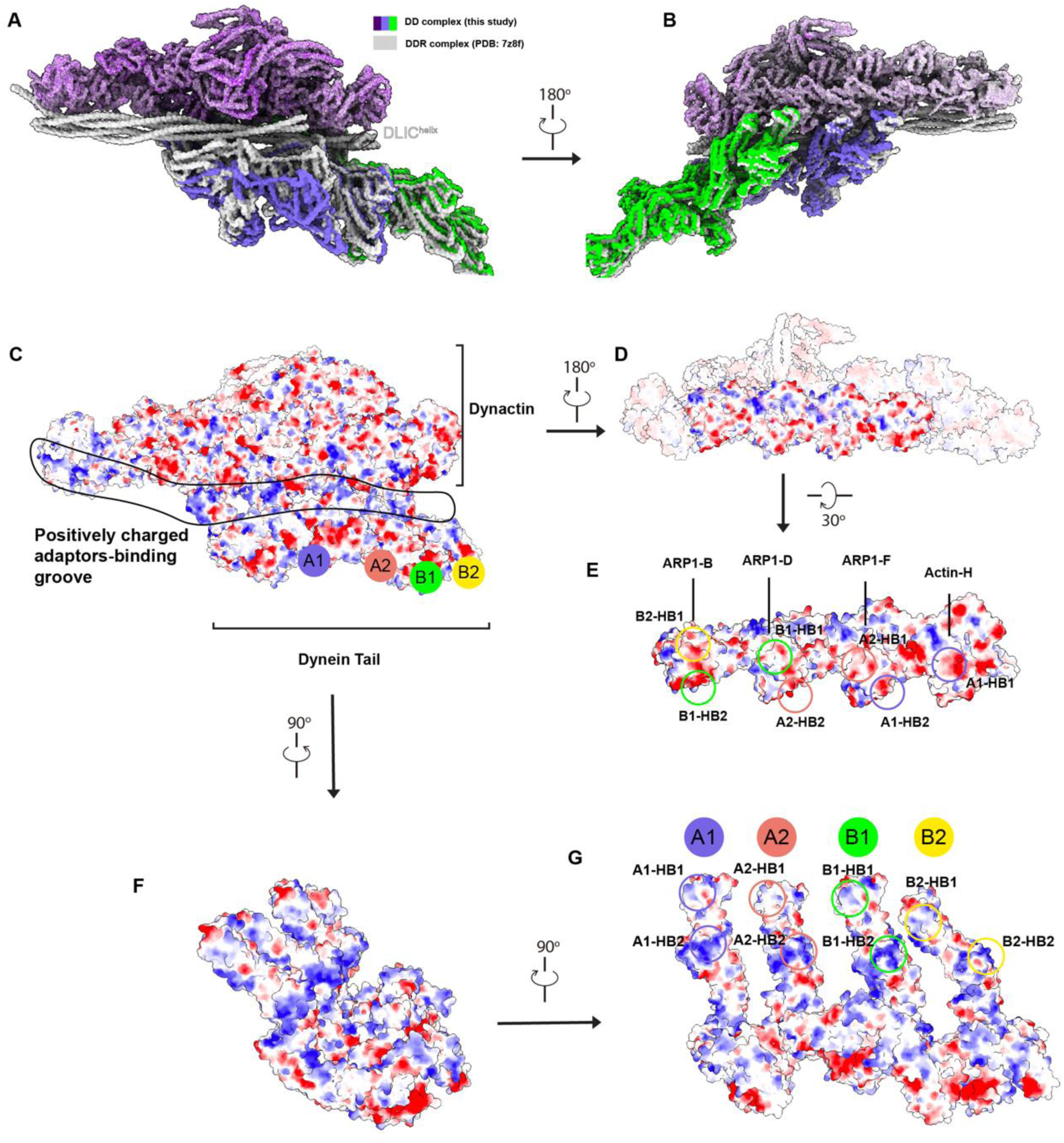
Dynein tail and dynactin interaction is facilitated by charge-charge interactions. (A-B) Superimposition of DD and DDR complex. The DLIC^helix^ is invisible in DD complex and is highlighted in DDR complex. (C) Surface electrostatics analysis of the dynein tail bound to dynactin, highlighting the positively charged adaptor-binding groove. (D-E) Two different views of electrostatic surface of the actin filament of dynactin. Circles indicate the interfaces corresponding to interactions with HB1/2 of the dynein tail in (G). (F-G) Two different views of electrostatic surface of the dynein tail. Circles indicate the interfaces corresponding to interactions with the dynactin actin filament in (E).

**Extended Data Figure 4.**
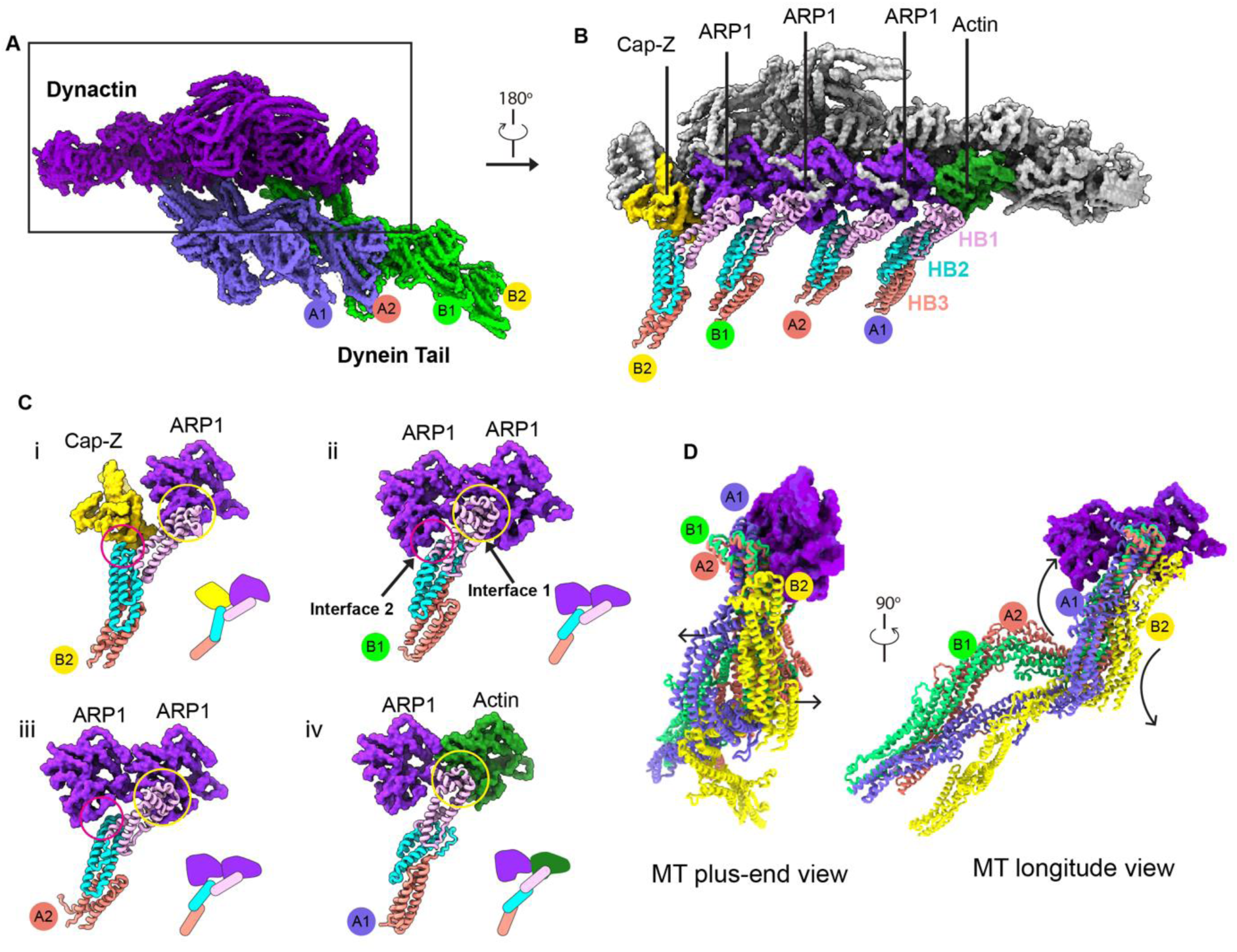
Interaction interfaces between dynein tail and dynactin actin filament part. **(A)** Molecular model of the dynein tail interacting with dynactin. **(B)** Opposite view shows the interactions between dynein tail HB1/HB2 and ARP1/Actin. For simplicity, NDD domain of dynein tail has been eliminated. **(C)** Detailed interfaces of each heavy chain interacting with dynactin, interface 1 and 2, along with cartoon models showing the interfaces. **(D)** Structural alignment of four dynein tails bound to the actin filament, arrows indicate the movement of heavy chains A1 and B2 relative to two middles aligned heavy chains A2/B1.

**Extended Data Figure 5.**
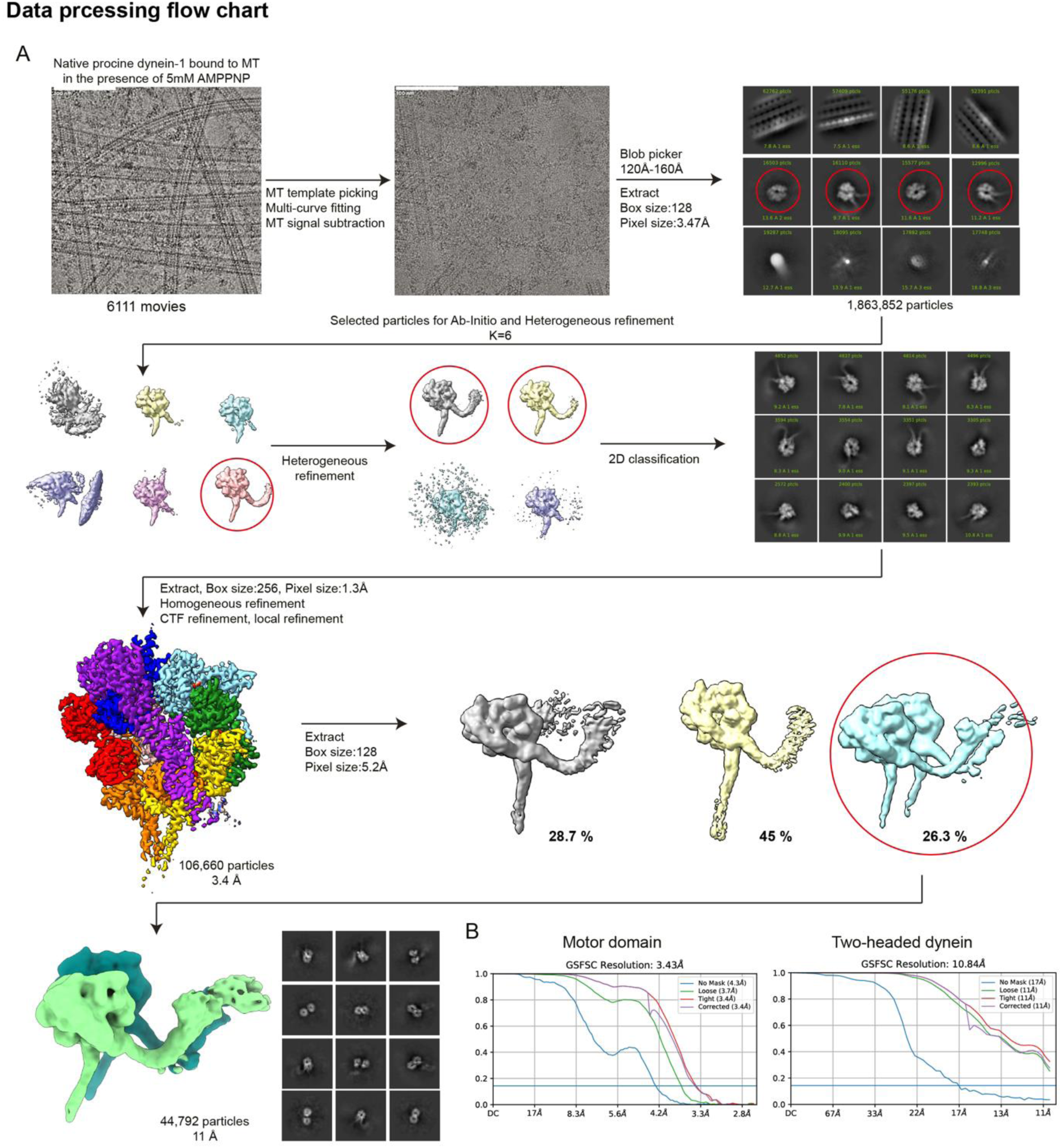
Data processing flow chart of D-MT. **(A)** Representative image of dynein bound to MTs and workflow of cryo-EM image processing. **(B)** FSC curves of motor domain and two-headed dynein reconstruction.

**Extended Data Figure 6.**
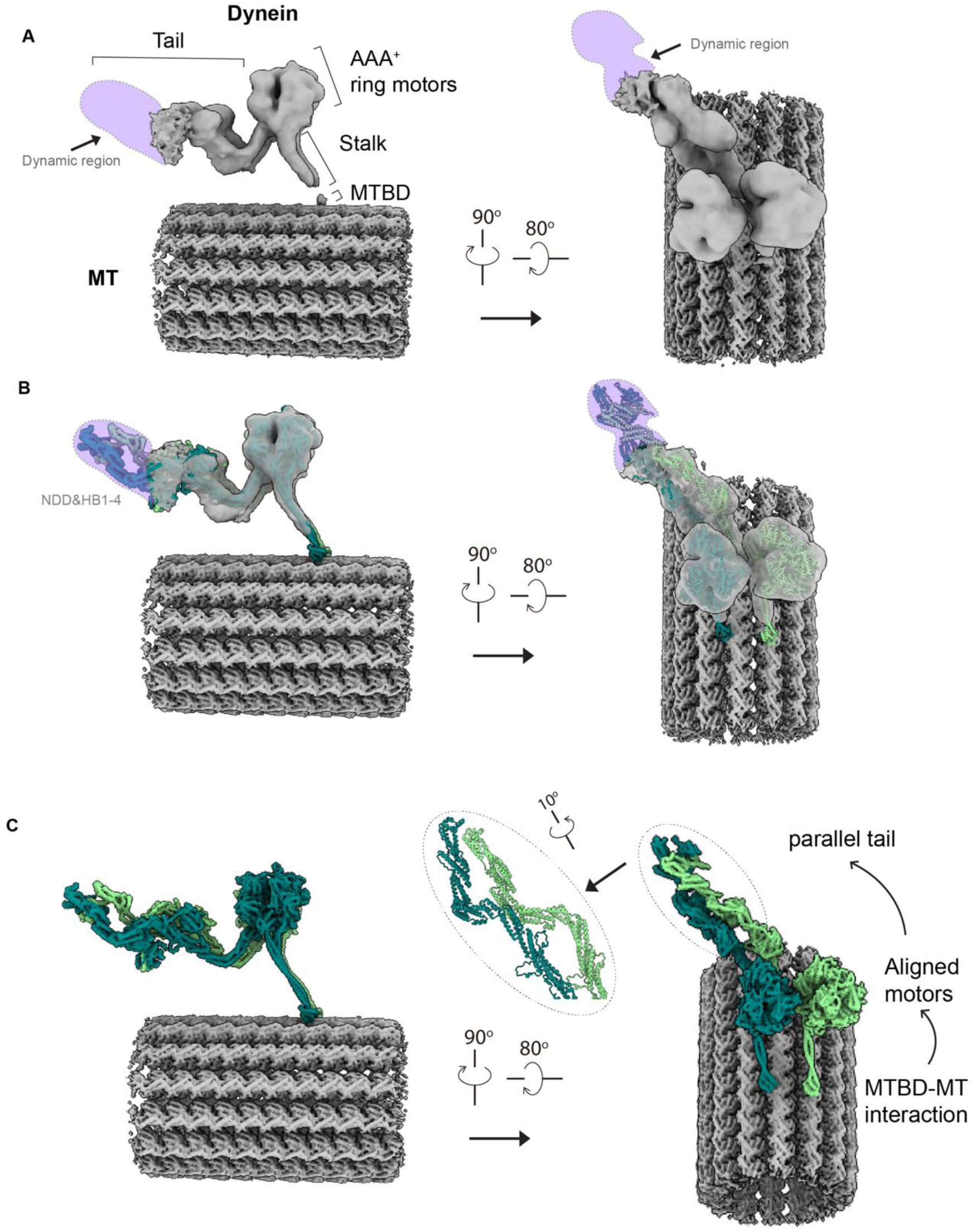
Cryo-EM structure of D-MT with two aligned motors. **(A)** Two different views of the density map of the D-MT complex are shown (44, 792 particles). The side view shows the dynein tail with a dynamic region, the aligned AAA+ ring motors, the stalk, and the microtubule-binding domain (MTBD) of dynein. The dynamic region of the tail is highlighted in light purple. **(B)** A molecular model of dynein-B, extracted from the DDR-MT complex (PDB: 7Z8F), is fitted into the cryo-EM structure of the D-MT complex (A) using rigid-body fitting. The NDD and HB1-4 domains of the dynein heavy chain contribute to the dynamic region. **(C)** The MT is displayed as a density map, while dynein is represented as a molecular model. A top view shows the parallel tails and the direction of conformational signal transmission, as indicated by arrows, upon interaction between the MTBD and MT.

**Extended Data Figure 7.**
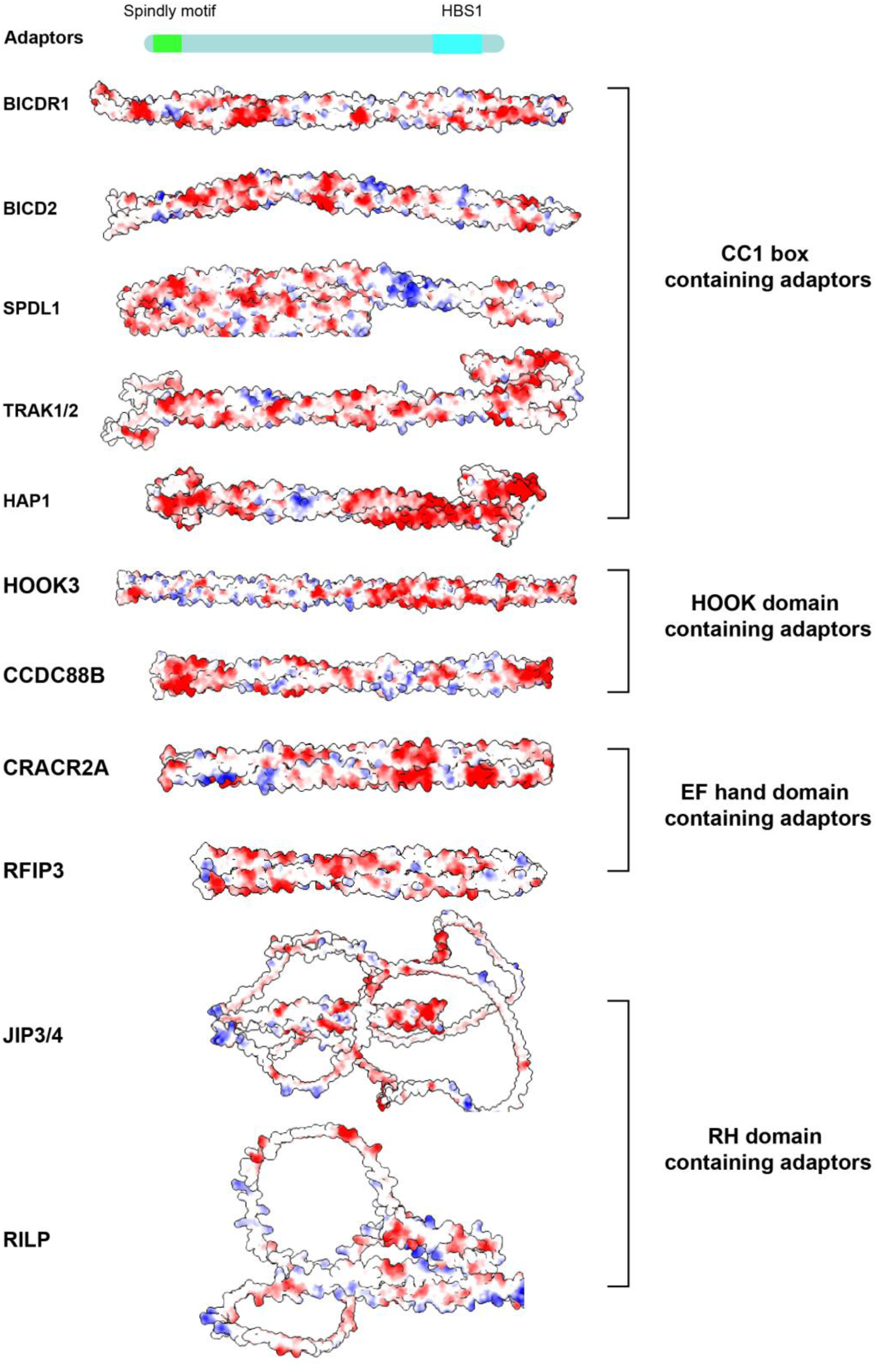
Surface electrostatics analysis of adaptors with domains corresponding to groove binding. The figure shows the surface electrostatics of various adaptors, categorized by their domain types. CC1 box containing adaptors: BICDR1, BICD2, SPDL1, TRAK1/2, HAP1. HOOK domain containing adaptors: HOOK3, CCDC88B. EF hand domain containing adaptors: CRACR2A, RFIP3. RH domain containing adaptors: JIP3/4, RILP

**Extended Data Figure 8.**
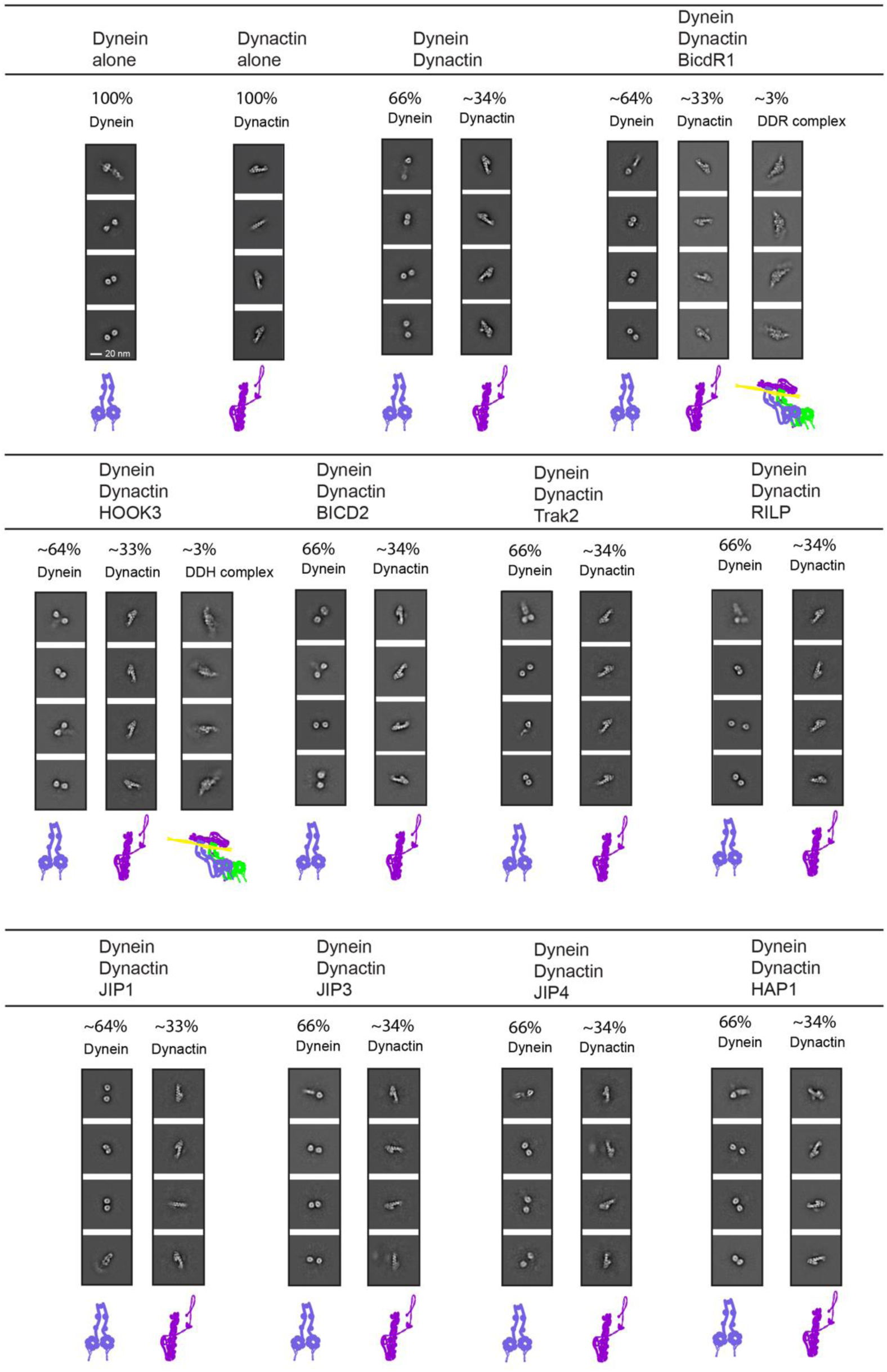
Negative-staining EM analysis of dynein, dynactin, dynein-dynactin complex, and dynein- dynactin-adaptor complexes. Representative 2D images of each sample and their corresponding populations are shown. The percentages indicate the proportion of dynein, dynactin, and dynein- dynactin-adaptor complexes observed in each condition. Scale bar, 20 nm.

**Extended Data Figure 9.**
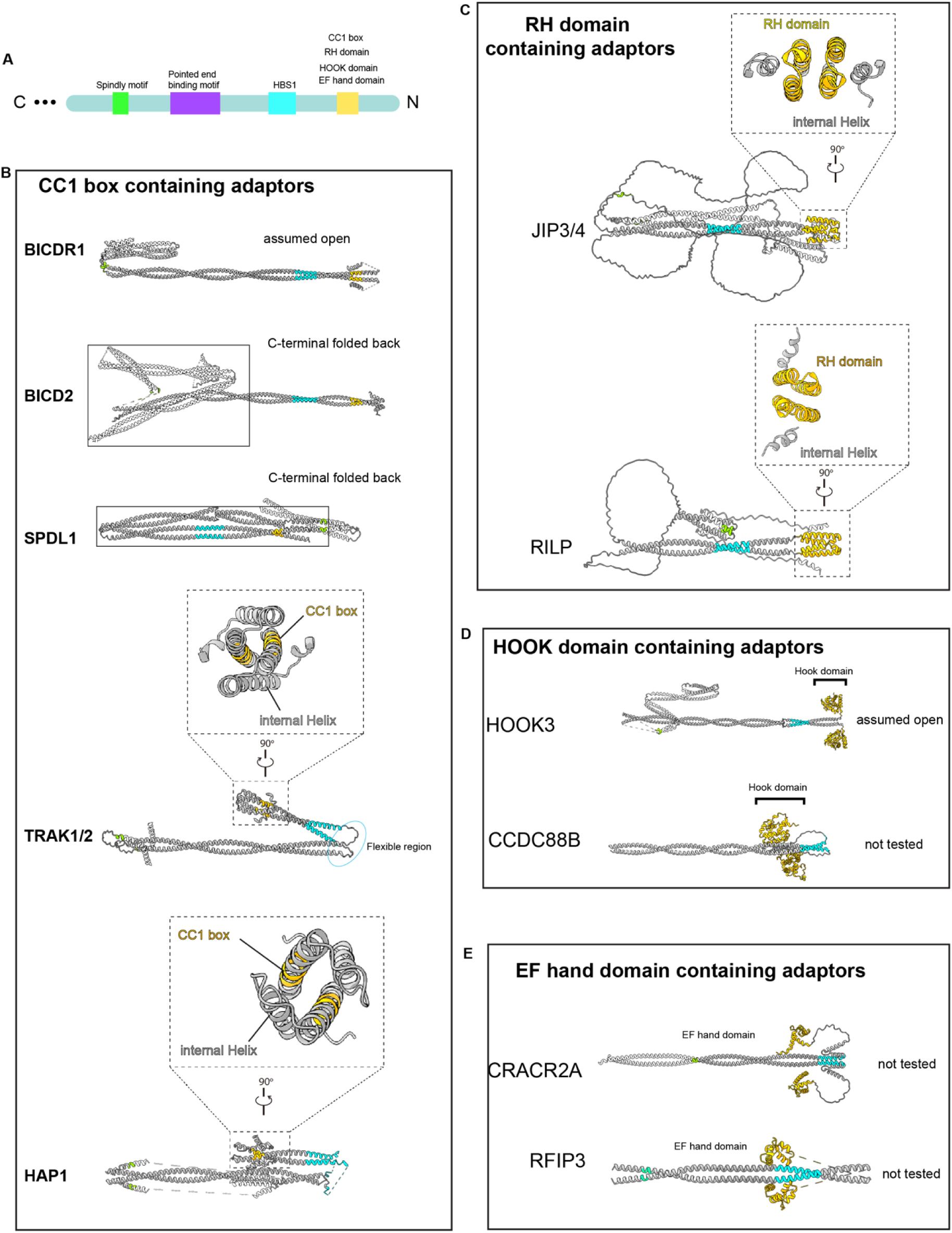
Schematics of adaptors and their AlphaFold2 predicted structures. **(A)** Domain structure of adaptors, showing the locations of the spindly motif, pointed end binding motif, HBS1, CC1 box, RH domain, HOOK domain, and EF hand domain. **(B)** CC1 box containing adaptors: BICDR1 (assumed open), BICD2 (C-terminal folded back), SPDL1 (C-terminal folded back), TRAK1/2, and HAP1. **(C)** RH domain containing adaptors: JIP3/4 and RILP. **(D)** HOOK domain containing adaptors: HOOK3 (assumed open) and CCDC88B. **(E)** EF hand domain containing adaptors: CRACR2A and RFIP3. Enlarged views of the autoinhibited structures of corresponding adaptors are enclosed in dashed boxes.

**Extended Data Figure 10.**
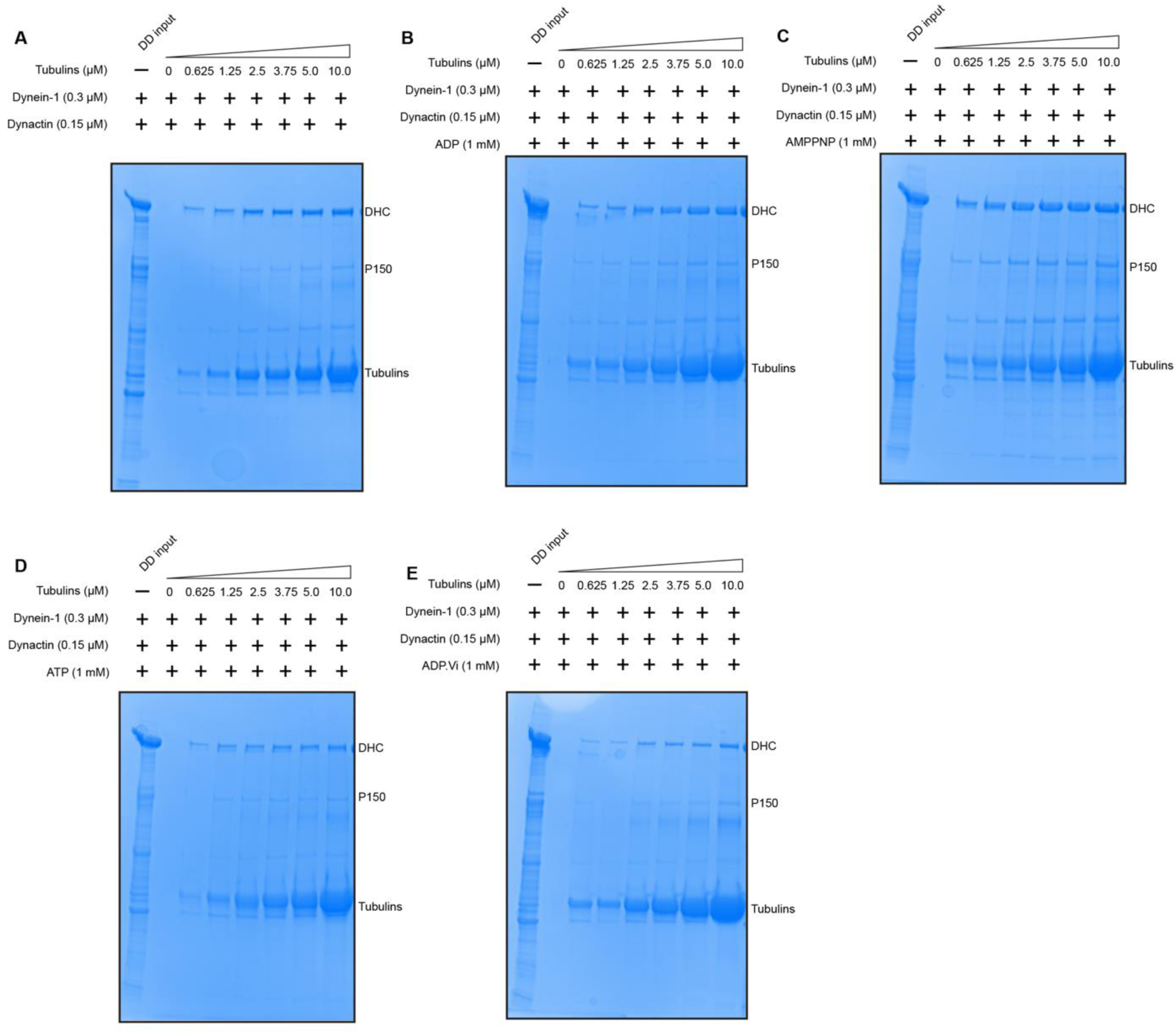
Binding affinity measurement of DD complex to MTs in different nucleotide binding states. SDS-PAGE gels of dynein-dynactin complex bound to MTs in various nucleotide binding states, **(A)** Apo, **(B)** ADP, **(C)** AMPPNP, **(D)** ATP, **(E)** ADP·Vi, stained with simple blue. All dynein samples used in this assay are in a nucleotide-free condition (Apo). Gels are representative of *n*=3 independent experiments.

**Extended Data Figure 11.**
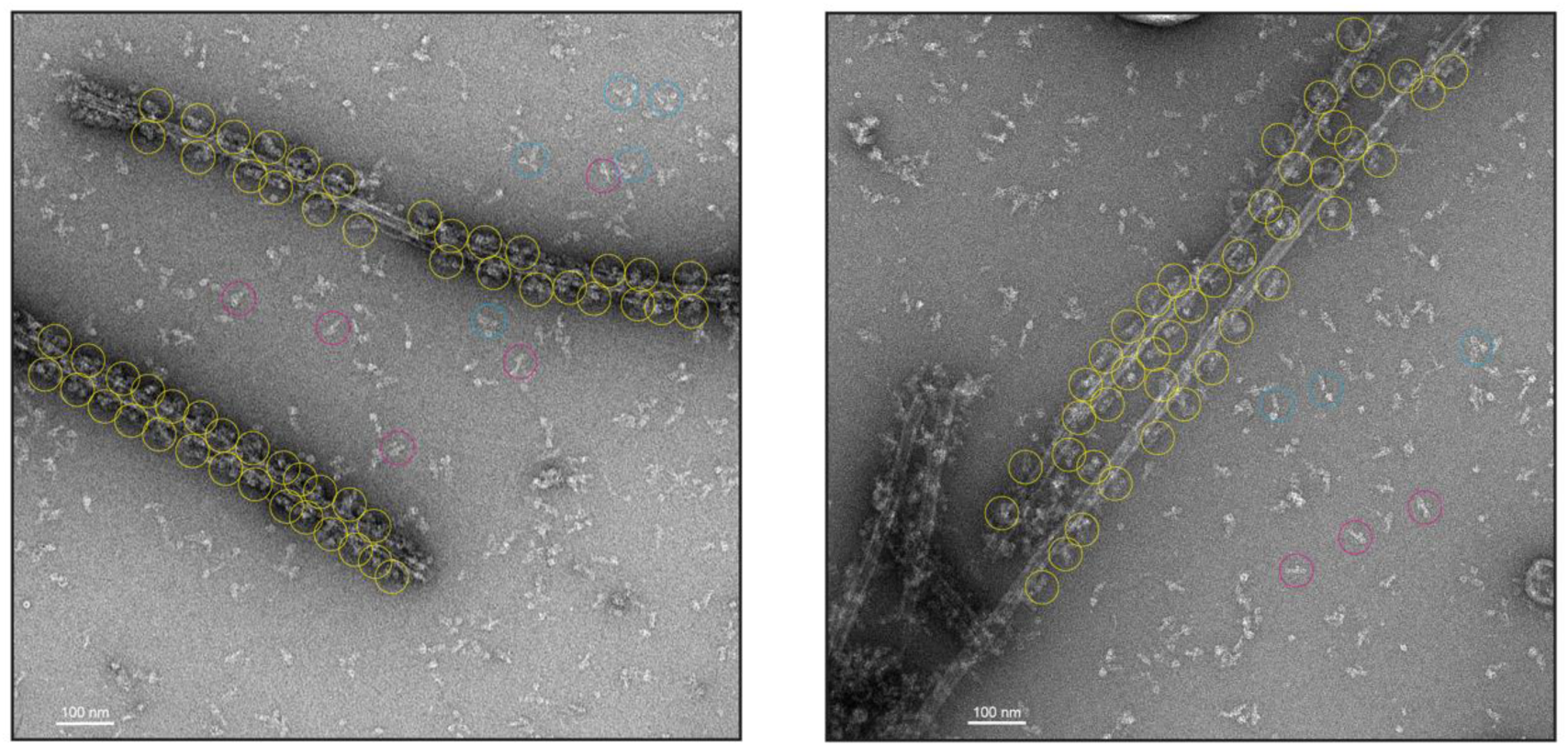
DDR complex enriched on MTs. Two representative negative-staining EM images (*n*=50) of the DDR-MT complex without any purification process. DDR complexes on MTs are manually identified and highlighted with yellow circles. Dynein (blue circles), dynactin (magenta circles), and DDR complexes off MTs were picked using cryoSPARC and subsequently subjected to 2D averaging analysis for identification. Scale bar: 100 nm.

**Extended Data Figure 12.**
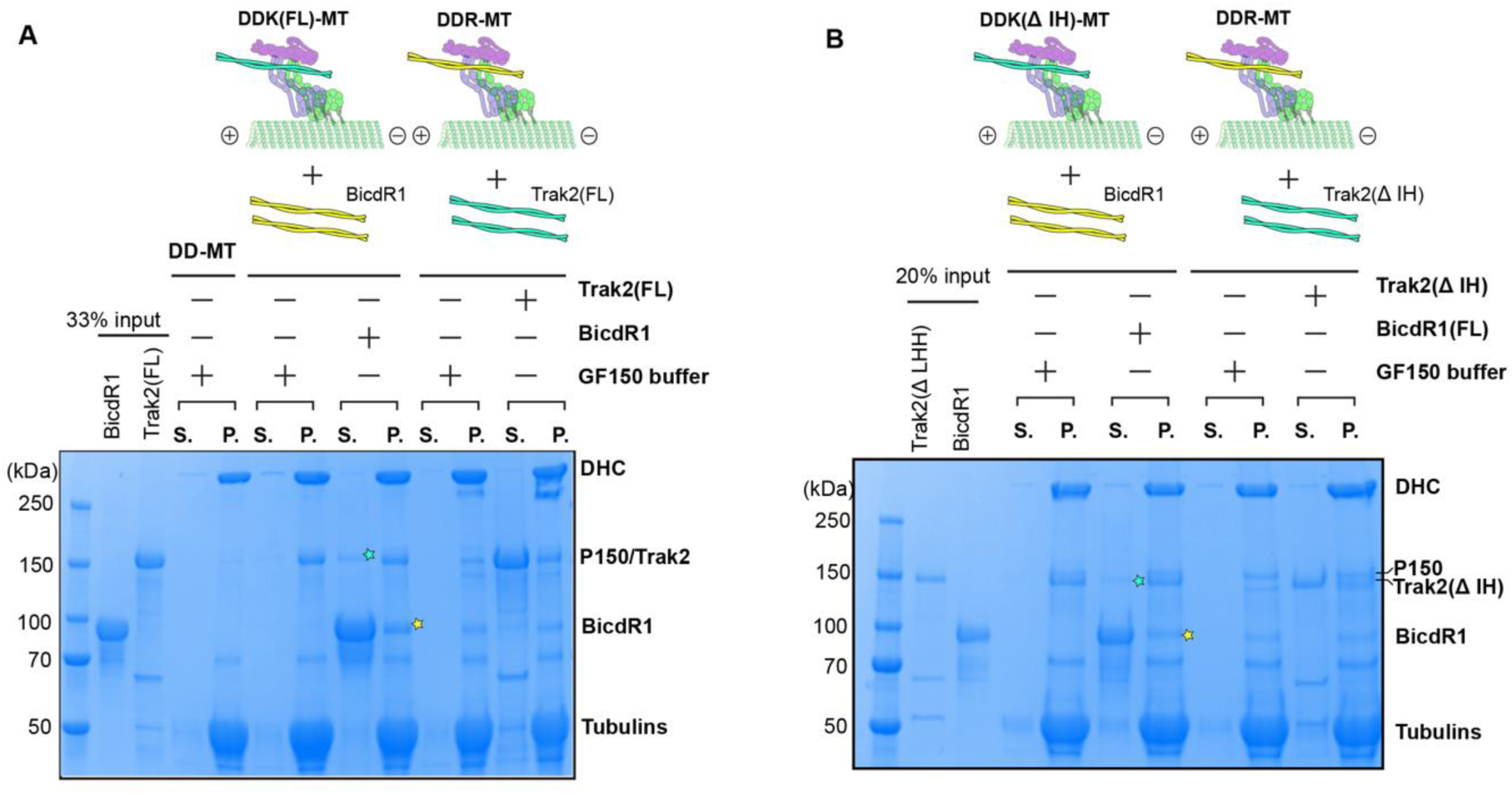
BicdR1 competes with Trak2 in pre-assembled DDK-MT complex. Schematic representation of the adaptor competition assay showing BicdR1 competing with the pre-formed DDK-MT and Trak2 competing with the pre-formed DDR complex. SDS-PAGE gels stained with simple blue show the results of these competition assays. Competition assay with full-length Trak2 (Trak2(FL) **(A)**, and truncated Trak2 (Trak2(Δ IH)) **(B)**. S, supernatant; P, pellet. The Trak2 competed off from the DDK-MT complex is marked with green stars, while the newly bound BicdR1 in the DDKR- MT complex is marked with yellow stars. Gels are representative of *n*=3 independent experiments.

**Extended Data Figure 13.**
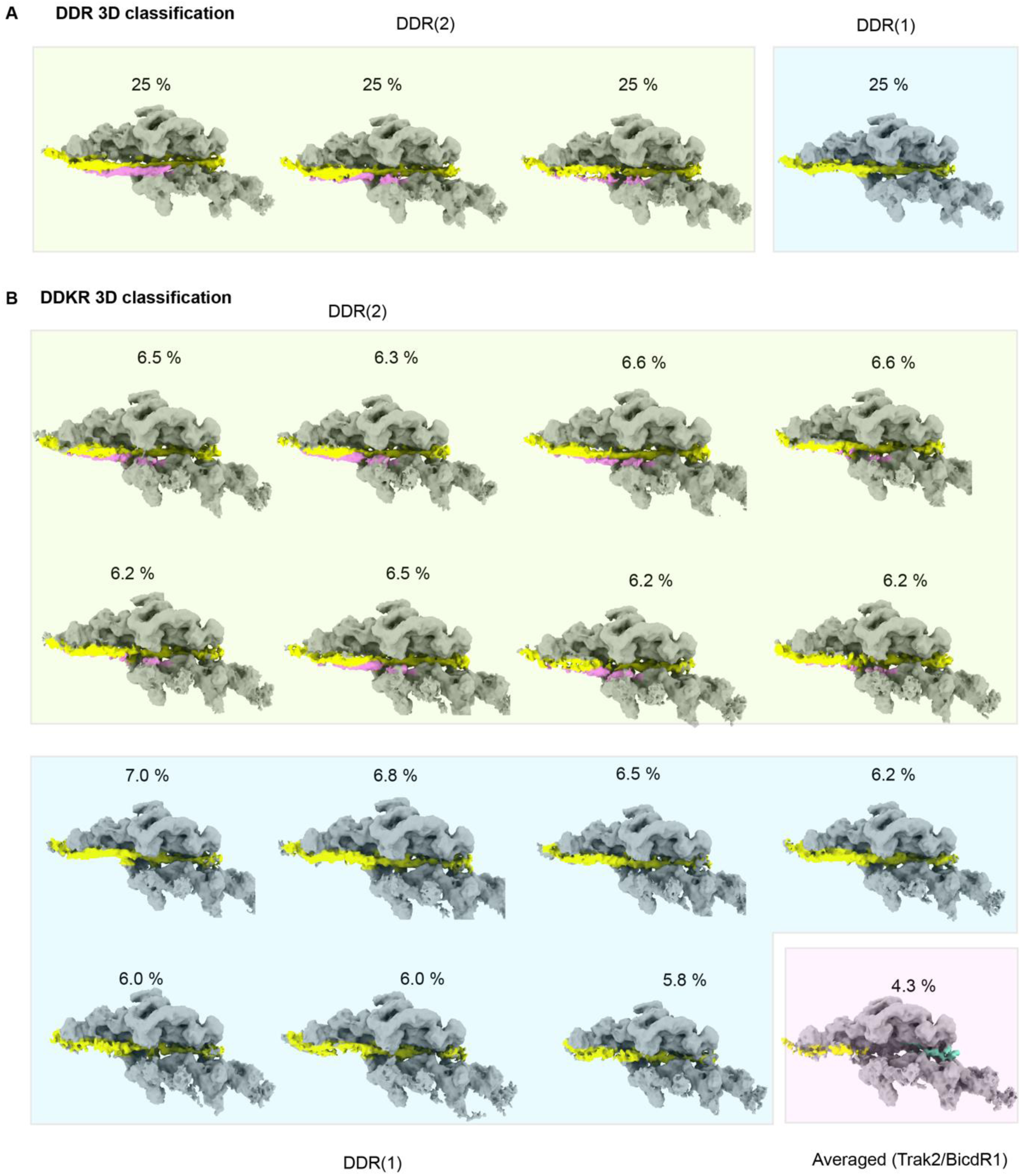
3D classification analysis of DDR-MT and DDKR-MT complexes. **(A)** 3D classification of DDR complexes showing two classes: DDR2 and DDR1, each representing 25% of the total particles. **(B)** 3D classification of DDKR complexes showing multiple subclasses of DDR2 and DDR1, with percentages indicating the proportion of each subclass. The averaged structure of DD- Trak2/BicdR1 is also shown, representing 4.3% of the total particles.

**Extended Data Table 1.** Cryo-EM data collection, refinement, and validation statistics.

| Description | Dynein-MT Overall | Dynein-MT Motor domain | Dynein-Dynactin-MT Overall | Dynein-Dynactin-MT Dynactin-Dynein-Tail | Dynein-Dynactin-BICDR1-MT One BICDR1 | Dynein-Dynactin-BICDR1-MT Two BICDR1 | Dynein-Dynactin-TRAK2-MT |
| --- | --- | --- | --- | --- | --- | --- | --- |
|  | EMD-46844 | EMD-46843 | EMD-46845 | EMD-46846 | EMD-46847 | EMD-46848 | EMD-46849 |
|  | PDB-9DGQ | PDB-9DGP | PDB-9DGR | PDB-9DGS | PDB-9DGT | PDB-9DGU | PDB-9DGV |
| **Data Collection and Processing** |  |  |  |  |  |  |  |
| Facility | Yale ScienceHill-Cryo-EM facility |  | Laboratory for BioMolecular Structure in BNL |  | Yale ScienceHill-Cryo-EM facility |  | BNL |
| Microscope | Glacios |  | Titan Krios |  | Glacios |  | Titan Krios |
| Voltage (kV) | 200 |  | 300 |  | 200 |  | 300 |
| Camera | K3 |  | K3 |  | K3 |  | K3 |
| Magnification | 45k |  | 105k |  | 45k |  | 105k |
| Pixel Size (Å) | 0.434 (super resolution) |  | 0.4125 (super resolution) |  | 0.434 (super resolution) |  | 0.4125 (super resolution) |
| Total Electron Exposure (e-/Å<sup>2</sup>) | 40 |  | 40 |  | 40 |  | 40 |
| Defocus Range (µm) | 1.5-2.7 |  | 1.5-2.7 |  | 1.5-2.7 |  | 1.5-2.7 |
| Symmetry Imposed | C1 | C1 | C1 | C1 | C1 | C1 | C1 |
| Num of mics | 6111 |  | 10660 |  | 3000 |  | 8369 |
| Initial Particles | 1 863 852 |  | 583 852 |  | 390 773 |  | 924 770 |
| Final Particles | 44 792 | 106 660 | 16 372 | 62 404 | 10 750 | 10 576 | 52 920 |
| **Refinement** |  |  |  |  |  |  |  |
| Initial models | 7z8f | 7z8f | 7z8f, 6znl | 7z8f, 6znl | 7z8f, 6znl | 7z8f, 6znl | 7z8f, 6znl |
| Map pixel size | 5.2 | 1.302 | 3.3 | 1.23 | 1.736 | 1.736 | 3.3 |
| Map Resolution (Å) (FSC 0.143) | 10.8 | 3.4 | 15 | 3.9 | 7.2 | 7.1 | 8.8 |
| Map sharpening B-factor (Å<sup>2</sup>) | NA | 66 | NA | 43.1 | NA | NA | NA |
| **Model Composition** |  |  |  |  |  |  |  |
| Non-hydrogen atoms | 32970 | 24594 | 131722 | 85495 | 89407 | 93353 | 86558 |
| Protein residues | 8153 | 3038 | 28258 | 11586 | 12054 | 12537 | 11800 |
| Ligands | MG: 2<br>ANP: 4<br>ADP: 2<br>ATP: 2 | MG: 4<br>ANP: 2<br>ADP: 1<br>ATP: 1 | MG: 1<br>ZN: 3 | ADP: 9<br>ATP: 1<br>ZN: 3 | ADP: 9<br>ATP: 1<br>ZN: 3 | ADP: 9<br>ATP: 1<br>ZN: 3 | ADP: 9<br>ATP: 1<br>ZN: 3 |
| **Model vs. Data** |  |  |  |  |  |  |  |
| FSC Map to Model (Å) (FSC 0.5) | NA | 3.7 | NA | 6.4 | NA | NA | NA |
| Correlation coefficient (mask) | NA | 0.77 | NA | 0.78 | 0.74 | 0.72 | 0.61 |
| **B factors (Å<sup>2</sup>)** |  |  |  |  |  |  |  |
| Protein | 76.33 | 68.82 | 464.7 | 300.6 | 328.4 | 348.55 | 308.78 |
| Ligand | 51.63 | 39.34 | 196.63 | 211.5 | 211.5 | 211.5 | 211.5 |
| **R.m.s deviation** |  |  |  |  |  |  |  |
| Bond length (Å) | 0.01 | 0.007 | 0.007 | 0.004 | 0.004 | 0.005 | 0.003 |
| Bond angles (°) | 1.513 | 1.022 | 1.309 | 0.97 | 0.701 | 0.983 | 0.63 |
| **Validation** |  |  |  |  |  |  |  |
| Molprobity score | 0.87 | 1.53 | 1.95 | 1.95 | 2.04 | 2.06 | 1.95 |
| Clashscore | 0.98 | 10.25 | 10.52 | 16.02 | 19.7 | 21.59 | 16.21 |
| Rotamer outliers (%) | 0 | 0.11 | 0 | 0.14 | 0.27 | 0.18 | 0.14 |
| **Ramachandran plot** |  |  |  |  |  |  |  |
| Outliers (%) | 0.02 | 0 | 0.4 | 0.1 | 0.1 | 0.1 | 0.12 |
| Allowed (%) | 2.33 | 1.33 | 5.78 | 3.56 | 3.6 | 3.4 | 3.51 |
| Favored (%) | 97.65 | 98.67 | 93.82 | 96.34 | 96.29 | 96.5 | 96.37 |
| **Rama-Z (whole)** | 1.21 | 0.91 | -0.83 | 0.53 | 0.35 | 0.29 | 0.52 |

